# Altered patterning of trisomy 21 interneuron progenitors

**DOI:** 10.1101/2020.02.18.951756

**Authors:** Yathindar Giffin-Rao, Jie Sheng, Bennett Strand, Ke Xu, Leslie Huang, Margaret Medo, Kirstin A. Risgaard, Samuel Dantinne, Sruti Mohan, Aratrika Keshan, Roger A. Daley, Bradley Levesque, Lindsey Amundson, Rebecca Reese, André M.M. Sousa, Yunlong Tao, Daifeng Wang, Su-Chun Zhang, Anita Bhattacharyya

## Abstract

Individuals with Down syndrome (DS, Ts21), the most common genetic cause of intellectual disability, have smaller brains that reflect fewer neurons at pre- and post-natal stages, implicating impaired neurogenesis during development. Our stereological analysis of adult DS cortex indicates a reduction of calretinin expressing interneurons. Using Ts21 human induced pluripotent stem cells (iPSCs) and isogenic controls, we find Ts21 progenitors generate fewer COUP-TFII+ progenitors with reduced proliferation. Single cell RNA-sequencing of Ts21 progenitors confirms the altered specification of progenitor subpopulations and identifies reduced WNT signaling. Activation of WNT signaling partially restores the COUP-TFII+ progenitor population in Ts21, suggesting that altered WNT signaling contributes to the defective development of cortical interneurons in DS.

## INTRODUCTION

The human cortex has evolved larger superficial layers and greater reliance on interneurons for complex functions than other mammals (Arshad et al., 2016; Dzaja et al., 2014; Hansen et al., 2013; Jones, 2009; Paredes et al., 2016; Radonjic et al., 2014). Defects in cortical interneuron development and function are linked to many neuropsychiatric disorders and disorders characterized by intellectual disability (Marin, 2012; Rossignol, 2011). Maldevelopment can lead to abnormal numbers, subtypes and/or placement of interneurons that significantly affect the functioning of the cortex, leading to cognitive impairment. The most common of these is Down syndrome (DS, Ts21), a complex multigene disorder caused by trisomy 21.

DS individuals have smaller brains (Becker et al., 1991; Davidoff, 1928; Schmidt-Sidor et al., 1990; Wisniewski, 1990) with reduced volume of frontal and temporal areas of the cortex, including the hippocampus (Emerson et al., 1995; Kesslak et al., 1994; Wisniewski, 1990). Morphological analysis of pre- and postnatal DS brains over the last century has consistently revealed fewer neurons in DS brain (Becker et al., 1991; Benda, 1947; Colon, 1972; Davidoff, 1928; Golden and Hyman, 1994; Ross et al., 1984; Takashima et al., 1981; Wisniewski et al., 1984), underlying the reduced volume and implicating reduced neurogenesis as a feature in DS (Lott and Dierssen, 2010). In particular, fewer granular cells and aspinous stellate interneurons are present in upper cortical layers in early life (Ross et al., 1984). These neuropathological studies were based on small sample sizes and morphological identification of interneurons. Our own *in vitro* data on DS fetal tissue corroborates that neuron reductions include GABAergic neurons (Bhattacharyya et al., 2009). We and others have modeled interneuron development using Ts21 induced pluripotent stem cells (iPSCs) and found that generation of interneurons from Ts21 iPSCs is altered, both *in vitro* and when transplanted *in vivo* (Huo et al., 2018; Xu et al., 2019). These observations point to the simple fact that the development of cortical interneurons is impaired in DS. The processes underlying the maldevelopment of cortical interneurons in DS remain unknown.

We address gaps in knowledge of interneuron development in DS through rigorous analysis of postmortem tissue to assess specific interneuron subpopulations for deficits and then model interneuron deficits using iPSCs from DS individuals and isogenic controls to reveal if and how the initial events of specification and proliferation of cortical interneurons are altered in DS. Our results show specific deficits in calretinin (CR+) interneurons and altered specification and proliferation of progenitors at the cellular level. Molecular data suggest reduced WNT signaling may be one of the mechanisms of faulty human interneuron development in DS.

## RESULTS

### Fewer interneurons in DS cortex

Limited histopathological observations reveal the presence of fewer neurons, primarily in superficial layers, in the cortex in DS (Golden and Hyman, 1994; Ross et al., 1984; Weitzdoerfer et al., 2001). Based on morphology, results suggest fewer aspinous stellate interneurons (Ross et al., 1984). We sought to determine whether there is a reduction in interneurons, and if so, to define which interneuron subtypes are altered, using immunohistochemical markers in DS cortex. We analyzed post-mortem superior temporal gyrus (STG) from four DS and four age-matched control male individuals (ages 15-35) (**Figure 1A, Table S1)**. These adolescent and young adult ages are similar to those in the previous study (Ross et al., 1984) and were chosen to avoid the early onset degeneration that occurs in DS as early as age 30 (Hartley et al., 2014). We analyzed the STG for several reasons: 1) thin STG is a consistent gross abnormality in DS brain (Becker, 1991; Mito et al., 1991); 2) alterations in neuron density and lamination in the STG occur during DS cortical development (Golden and Hyman, 1994); and 3) the STG is an association cortex likely to be involved in intellectual impairment in DS. Immunocytochemistry and design-based stereology were used to quantify neuronal populations in sections (Perl et al., 2000; West, 2013) to ensure unbiased, efficient, and more reliable results than other ad hoc quantitative analyses (Boyce et al., 2010) and to take into account the gross size differences of DS brains. We used the Optical Fractionator Probe method (Stereo Investigator, MBF) to uniformly sample and estimate cell number and the Cavalieri probe method to estimate volume (West et al., 1991). Our results show a reduced density of NeuN+ cells, corroborating the reduction in the number of neurons in DS cortex (**Figure 1B**; Control = 8.33 x 10^6^ ± 0.79 x 10^6^, DS= 4.90 x 10^6^ ± 0.15 x 10^6^, p=0.029, N=4). Using parvalbumin (PV) to identify the predominant interneuron subtype in mouse, we observed no density difference in this population between DS and controls (Control = 0.98 x 10^6^ ± 0.18 x 10^6^, DS= 0.89 x 10^6^ ± 0.24 x 10^6^, p=0.89, N=4). Using CR as a marker of upper layer interneurons, we find a greater proportion of CR+ neurons than PV+ neurons in both control and DS STG samples. The density of CR+ cells may be reduced in DS samples (Control = 1.74 x 10^6^ ± 0.14 x 10^6^, DS= 1.19 x 10^6^ ± 0.23 x 10^6^, *p=0.11,* N=4), although a larger sample size is needed to reach statistical significance. To assess whether neuron counts correlate with post-mortem interval (PMI), independent of genotype, we performed a Pearson correlation analysis. The results show that there is no relationship between PMI and PV (0.05), but there is a moderate relationship between PMI and NeuN (−0.41) and CR (−0.59). Despite these technical considerations, the results identify differences in the presence of interneuron subtypes, CR+ but not PV+, in adult DS cortex.

**Figure 1:**
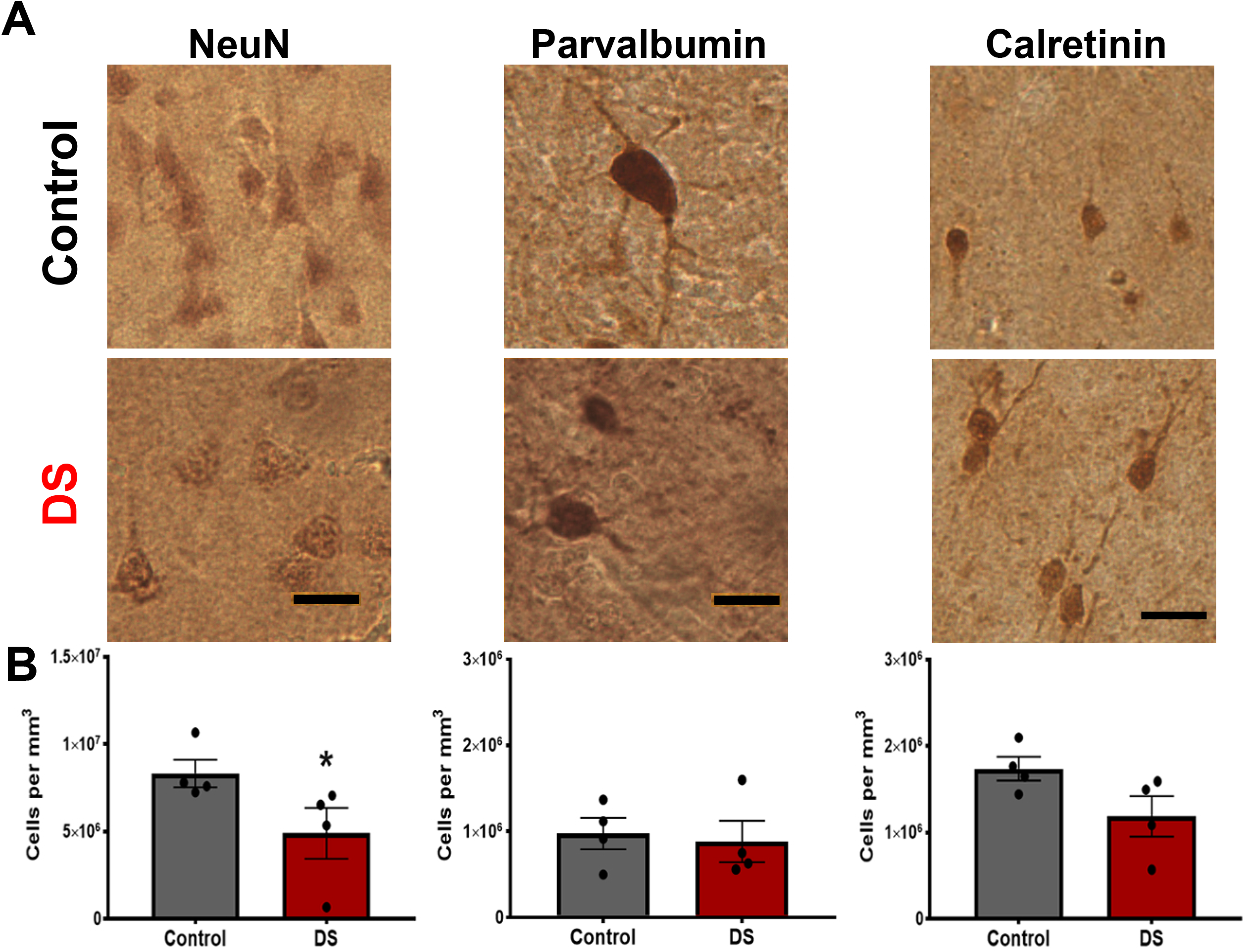
Reduced density of neurons in post-mortem Down syndrome cortex. A) Representative images of NeuN+, PV+, and CR+ neurons detected by immunohistochemistry (DAB) in control and DS post-mortem superior temporal gyrus (STG) samples. B) Stereological quantification of the density of each neuron subtype neurons in control and DS tissue. (N=4; NeuN *p=0.03, PV p=0.89, CR p=0.11 calculated using non-parametric Mann-Whitney U test).

### Ts21 iPSCs generate fewer CR+ neurons and fewer COUP-TFII+ progenitors

Our *in vivo* results corroborate historical data and suggest that the generation of cortical interneurons is altered by Ts21. Interneuron progenitors are specified and proliferate in the three neurogenic areas of the ventral telencephalon: the lateral, medial, and caudal ganglionic eminences (LGE, MGE and CGE) (Anderson et al., 2002; Flames and Marin, 2005; Kessaris et al., 2014; Parnavelas et al., 2000; Xu et al., 2004). Individual spatially-defined progenitor subpopulations with distinct transcriptional profiles each ultimately give rise to specific interneuron subtypes, thus initiating the diversity of interneurons in the cortex. MGE progenitors express the transcription factor NKX2.1 and are the primary source of interneurons in mouse, giving rise to 70% of the total cortical interneurons including PV, calbindin and somatostatin expressing subtypes (Du et al., 2008; Xu et al., 2010; Xu et al., 2008). CGE progenitors express the transcription factor COUP-TFII/NR2F2 that is also expressed by progenitors in the caudal MGE and along the MGE/LGE boundary (Kanatani et al., 2008). The CGE gives rise to vasoactive intestinal polypeptide, CR, reelin and neuropeptide Y expressing interneurons in mouse (Xu et al., 2004). Recent work indicates that these transcription factor codes are largely conserved between mouse and human, but human interneuron progenitors are more complex (Shi et al., 2021).

Here, we explore the development of human interneuron progenitors using iPSC modeling. Strategies to generate ventral neural progenitor cells (NPCs) and cortical interneurons from hPSCs (Kim et al., 2014; Liu et al., 2013; Maroof et al., 2013; Nicholas et al., 2013) rely on exogenous addition of sonic hedgehog (SHH) to specify progenitors to a ventral fate and yield a highly enriched population of GABA interneurons that mature to specific interneuron subtypes. One pair of isogenic Ts21 and control iPSCs (WC-24) were differentiated to cortical interneuron progenitors with exogenous sonic hedgehog (SHH) from Day 10-17 (**Figure 2A, Table S2**) (Liu et al., 2013). We previously reported the reduced generation of CR+ neurons from Ts21 progenitors using this paradigm (Huo et al., 2018). We repeated the experiment by maintaining our isogenic control and Ts21 interneuron progenitors for 74 or 91 days and then differentiating them to neurons to assess the generation of CR+ neurons. Quantification of CR+ neurons reveals reduced generation of CR+ neurons from Ts21 progenitors (**Figure 2B**), corroborating our previous results in distinct cell lines and confirming we can model the decreased population of CR+ neurons in DS that we observe *in vivo* (**Figure 1**). Interneuron progenitors are specified temporally in addition to the spatial patterning that we are modeling in our culture system and expression of subtype-specific markers *in vitro* appears after long-term culture and so we acknowledge that when counting at a particular timepoint, the number does not necessarily equal the progenitor population.

**Figure 2:**
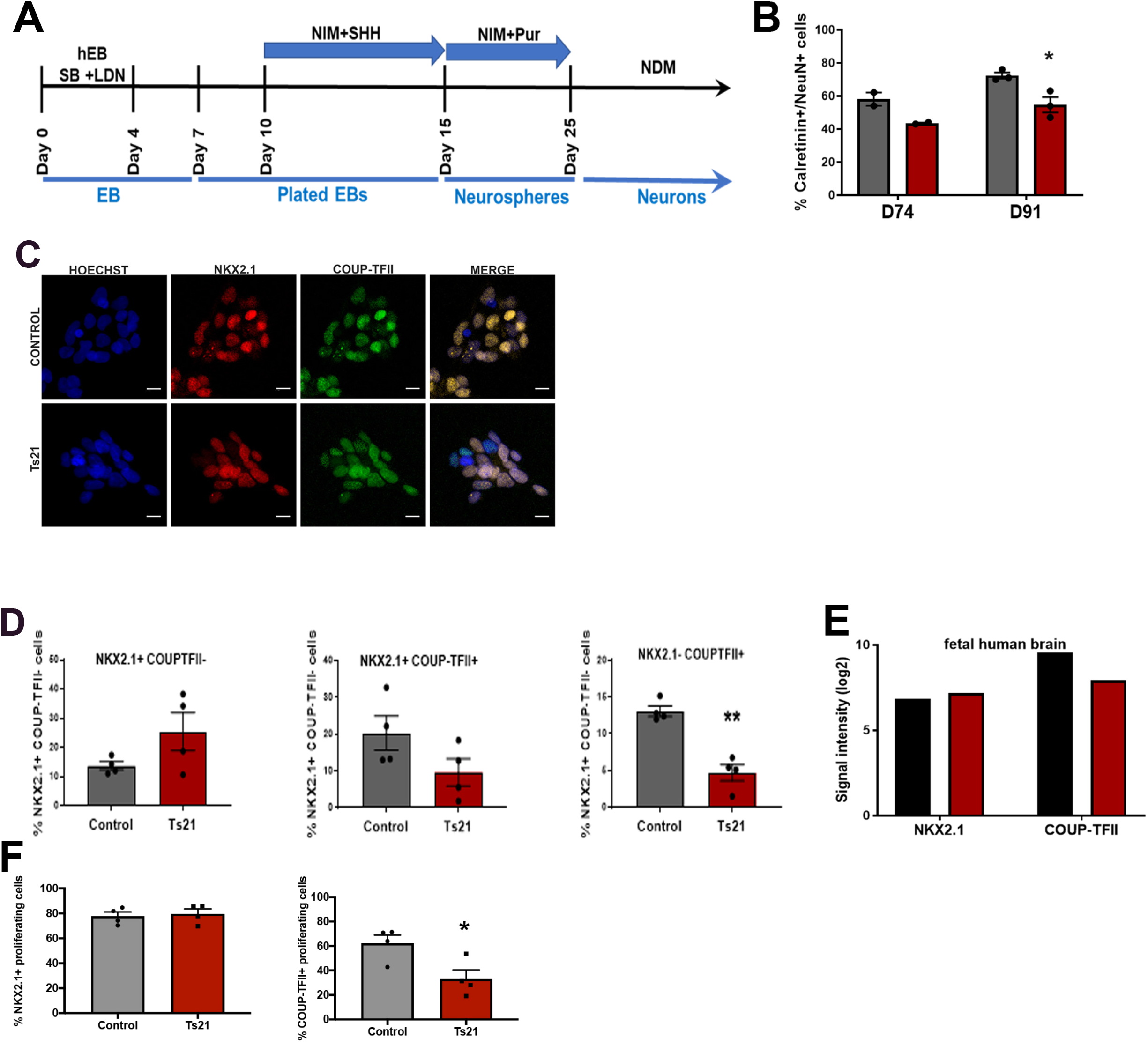
Ts21 iPSCs generate fewer fewer CR+ interneurons and fewer COUP-TFII+ progenitors. A) Interneuron progenitor differentiation protocol. B) Proportion of calretinin (CR) neurons/(NeuN+) differentiated from control and Ts21 progenitors at Day 74 and Day 91. *p= 0.023 using unpaired t-test with Two-stage step-up (Benjamini, Krieger, and Yekutieli). C) Immunofluorescence images of NKX2.1+ and COUP-TFII+ nuclei in control and Ts21 neural progenitor cells (NPCs). D) Proportion of NKX2.1+ COUP-TFII-cells (MGE); NKX2.1+ COUP-TFII+ cells (caudal MGE) and NKX2.1-COUP-TFII+ cells (CGE) in control and Ts21 NPCs. (**p-value <0.001 using unpaired t-test, N=4 cell lines). E) Expression of NKX2.1 and COUP-TFII in human fetal control and Down syndrome brain (14-17 weeks’ gestation, dorsolateral forebrain) (Olmos-Serrano et al., 2016). F) NKX2.1+/EdU+ and COUP-TFII+/EdU+ proliferating cells in the Ts21 cells compared to controls (*p-value<0.05 using unpaired t-test, N=4).

We focused on the specification of interneuron progenitors to gain insight into early developmental events that go awry in DS. NKX2.1+ and COUP-TFII+ populations are not mutually exclusive. While NKX2.1 is expressed in MGE, COUP-TFII is expressed in two spatial subpopulations of cortical interneuron progenitors: caudal MGE and CGE (Anderson, 2002; Anderson et al., 2002; Campbell, 2003; Parnavelas et al., 2000; Wilson and Rubenstein, 2000). These two COUP-TFII+ populations can be distinguished by NKX2.1 expression (Alfano et al., 2014; Kanatani et al., 2008; Lodato et al., 2011; Reinchisi et al., 2012); MGE progenitors express only NKX2.1, CGE progenitors express only COUP-TFII, while caudal MGE progenitors express both. We differentiated four Ts21 iPSC and control iPSC lines (**Table S2**) using our paradigm (**Figure 2A**). To test whether Ts21 affects specification of interneuron progenitors, we quantified the subpopulations of progenitors. Immunofluorescence of progenitors at Day 17 indicates that both control and Ts21 progenitors respond to SHH by expressing NKX2.1, as expected, as well as expressing COUP-TFII (**Figure 2C**). The percentage of MGE progenitors, as defined by NKX2.1+ COUP-TFII-expression, is not consistently different in Ts21 progenitors compared to isogenic or non-isogenic controls (Control = 13.65 ± 11.40, Ts21= 25.43 ± 6.49, p=0.1264, N=4) **(Figure 2D)**. The proportion of NKX2.1+COUP-TFII+ cells that represent caudal MGE-like cells is not statistically different between Ts21 iPSCs and controls (Control = 20.28 ± 4.64, Ts21= 9.632 ± 3.70, p=0.1232, N=4) **(Figure 2D)**. The low proportion of NKX2.1+ MGE progenitors overall is likely to the relatively late addition of SHH (at Day 10) in our protocol. Quantification of COUP-TFII+, NKX2.1-cells that represent CGE-like progenitors revealed fewer of these cells differentiate from Ts21 iPSCs compared to controls (Control = 13.09 ± 0.72, Ts21= 4.71 ± 1.12, p=0.0007, N=4) (**Figure 2D**). The results indicate that Ts21 progenitors generate fewer COUP-TFII+ cells.

Differences in expression of NKX2.1 and COUP-TFII expressing progenitors recapitulate *in vivo* development as there is decreased expression of *COUP-TFII/NR2F2*, but not *NKX2.1*, in human fetal DS brain (Olmos-Serrano et al., 2016) (**Figure 2E**). As COUP-TFII+ progenitors are an important source of CR interneurons in mice (Xu et al., 2004), the decreased generation of COUP-TFII+ progenitors from Ts21 iPSCs and reduced expression of *COUP-TFII/NR2F* in DS fetal brain provide a potential mechanism for the decreased population of CR+ interneurons in DS.

### Fewer Ts21 COUP-TFII+ progenitors are proliferating

Progenitors proliferate in response to extrinsic cues to expand the progenitor pool. We quantified proliferation of NPCs derived from three pairs of Ts21 and control iPSC lines by pulsing cells with EdU for 8 hours to label dividing cells. We asked if the proportions of proliferating NKX2.1+ and COUP-TFII+ cells differed between Ts21 and controls. Co-labeling of EdU+ cells with NKX2.1 or COUPTF-II in shows no difference in the proportion of dividing NKX2.1+ cells in Ts21 (**Figure 2F**), indicating that the NKX2.1+ population is preserved in Ts21. In contrast, fewer dividing COUP-TFII+ are found in the Ts21 population than controls (**Figure 2F**). These data suggest that Ts21 COUP-TFII+ cells are either proliferating more slowly or exiting the cell cycle prematurely.

### Single-cell transcriptomic analysis confirms cellular and molecular differences in Ts21 progenitors

To identify putative progenitor subpopulations that differ in Ts21 and to define gene pathways that are dysregulated by Ts21, we carried out single cell RNA-sequencing (scRNA-Seq) analysis using ventralized progenitors at Day 17 from one pair of isogenic Ts21 and control lines (WC24). Isogenic pairs of iPSCs are extremely valuable for molecular profiling where variation between individuals and cell lines can be amplified and mask subtle differences.

As we are investigating an actively proliferating population of cells and because there are differences in the proliferation of Ts21 progenitors, cell cycle genes are likely to be overrepresented in the analysis and therefore lead to clustering of cells based upon cell cycle rather than cellular fate. We used Seurat (Version 4.0.4) to regress out cell cycle genes so that the underlying gene markers could be used to segregate Ts21 and control cell populations (Butler et al., 2018; Mayer et al., 2018). The resulting UMAP plot shows the euploid and Ts21 progenitors overlaid on each other, revealing similar populations in Ts21 and controls (**Figure 3A**). Feature plots of the data indicate that that *NKX2-1* expression is restricted to a subset of clusters (**Figure 3B**). In contrast, *COUP-TFII/NR2F2* expression is widespread (**Figure 3C**), in agreement with widespread expression of *COUP-TFII/NR2F2* in fetal human ganglionic eminences (CGE and LGE) *in vivo* (Shi et al., 2021). The expression of these genes is thus different from protein expression (**Figure 2**).

**Figure 3:**
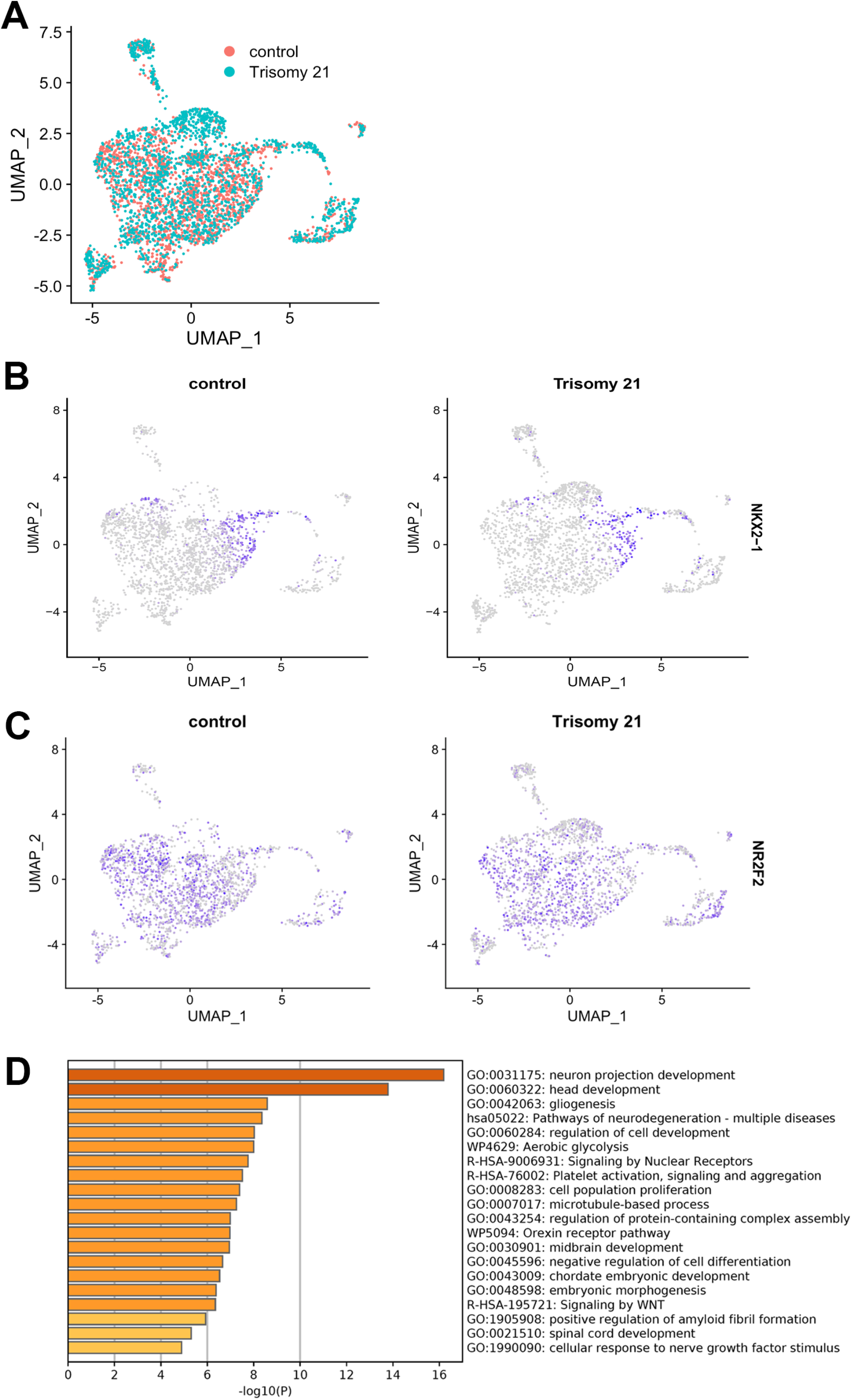
Single cell RNA-seq reveals gene expression differences in Ts21 progenitors. A) Dimensional reduction by experimental group (control and trisomy 21). B) Feature plot of expression of NKX2.1 in control and trisomy cells. C) Feature plot of expression of NKX2.1 in control and trisomy cells. D) Pathway analysis of differentially expressed genes in Ts21 compared to control.

Differentially expressed genes (DEGs) between Ts21 and control progenitor populations were identified using FindMarkers() function in Seurat with parameters logfc.threshold > 0.25 and padj value of <0.05 and 206 DEGs were identified (**Table S6**). Pathway analysis of DEGs between Ts21 and control using Metascape (Zhou et al., 2019) indicates genes involved in several neurodevelopmental processes are disrupted by Ts21 (**Figure 3D**, **Table S7, S8**). Association of DS and familial Alzheimer’s disease with our data was validated using the DisGeNet database (**Figure S1**). Pathways, including both neurodevelopmental and neurodegenerative pathways, are driven by the increased expression of the chromosome 21 encoded gene *APP* in the Ts21 progenitors. Neurodevelopmental pathways emerge due to the altered expression of key transcription factors, including *FOXG1, DLX2, LHX2, HES1*. WNT signaling emerges as an affected pathway driven by altered expression of non-chromosome 21 genes (*CALM2, SOX3, GNG2 and GNG3, RSPO3* and *RSPO1, RAC1, PFN1* and *TCFL2*). *RSPO1* and *RSPO3*, context dependent regulators of WNT signaling, are increased in Ts21 progenitors (Jin and Yoon, 2012; Rong et al., 2014) while *TCFL2,* a downstream effector gene (Chodelkova et al., 2018), is decreased. These single cell data identify WNT as a dysregulated pathway in Ts21 interneuron progenitors.

### Single-cell clustering reveals a GLI3 expressing subpopulation enriched in Ts21 progenitors

To identify putative progenitor subpopulations that differ in Ts21, we used Seurat to cluster the cells to reveal 15 subpopulations (**Figure 4A**). The identity of these cell populations was defined by identifying DEGs in each cluster with combined p-values less than 0.05 (**Table S9, Figure S2**). We cross-referenced these gene signatures to known gene markers of NPC populations to classify each population. The clustered cell types expressed genes indicative of cells at different stages of differentiation and different progenitor populations, revealing the heterogeneity of our iPSC-derived progenitors. We also compared cluster marker expression in our cells with clusters of human fetal ganglionic eminences (Shi et al., 2021) and found widespread expression overlap, validating that our progenitors have similar gene expression as fetal tissue (**Figure S3**). Similar proportions of control and Ts21 cells were represented in each subpopulation **(Figure 4B)**. However, quantification of the number of cells in each cluster revealed that one cluster, Cluster 3, was enriched in Ts21 **(Figure 4C)**. This cluster is identified by expression of *FOXG1* and the long noncoding RNA *LINC00551* **(Figure 4D,Table S9)**. Interestingly, *GLI3* is a unique marker gene for Cluster 3 (**Figure 4E, Table S9**). *GLI3*, as well as other marker genes in this cluster (*FOXG1, LHX2, MEIS2*) are highly expressed in intermediate progenitor cells in fetal brain (Li et al., 2018) and thus confirms the identity of Cluster 3 as a population of intermediate progenitors.

**Figure 4:**
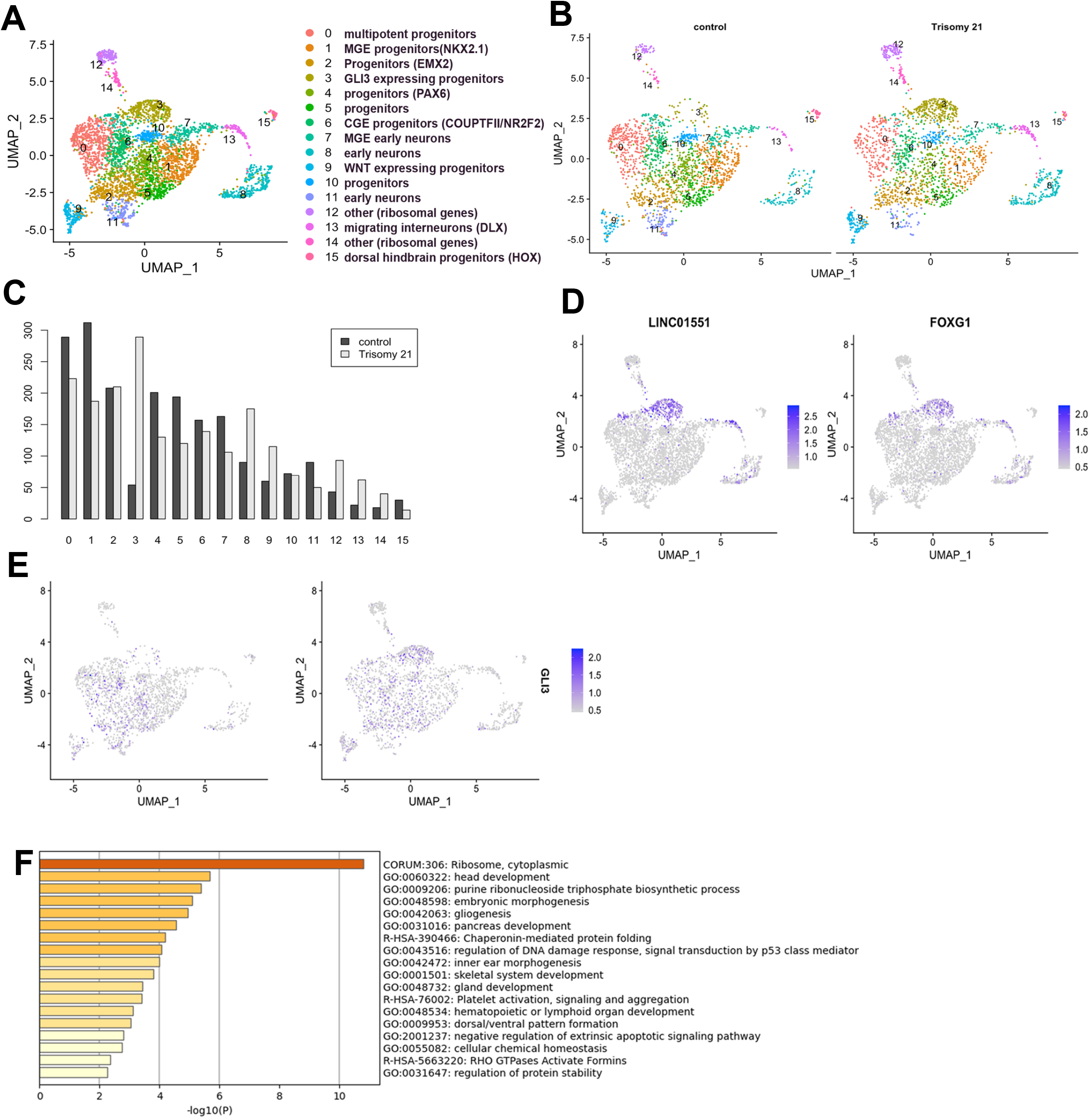
Single cell RNA-Seq reveals differences in Ts21 progenitor clustering. A) Clustering analysis revealed 15 sub-populations of cells based on expression of known gene markers. B) Clustering of control (2134 cells) and Ts21 (2158 cells). C) Proportion of control and Ts21 cells in each cluster reveals enrichment of Cluster 3 in Ts21 cells. D) Feature plot of LINC0551 and FOXG1, markers that identify Cluster 3 (Table S9). E) Feature plot showing expression of GLI3. F) Pathway analysis of differentially expressed genes in Cluster 3 in Ts21 compared to control.

DEGs between Ts21 and control cells in Cluster 3 were identified using FindMarkers() function in Seurat with parameters logfc.threshold > 0.25 and padj value of <0.05 and 72 DEGs were identified (**Table S10**). Pathway analysis of these DEGs using Metascape (Zhou et al., 2019) indicates genes involved in several cellular processes are disrupted by Ts21 (**Figure 4F, Table S11**). Enriched expression of ribosomal (*RPL37A, RPL4*) in Ts21 cells drives the identification of the top pathway of ribosome/translation. These genes are also enriched in intermediate progenitors (Li et al., 2018). The chromosome 21 gene *ATP5PF* drives the identification of the purine biosynthesis. DEGs in Cluster 3 include down regulation of WNT pathway genes in Ts21 (*TCF7L2, SOX4, GNG3* and *GNG5*, and *GPC3*). Together, these data raise the hypothesis that Cluster 3 arises from Ts21 progenitors due to decreased WNT signaling.

### Ts21 progenitors exhibit reduced WNT and upregulated GLI expression that can be rescued by WNT activation

Cellular analysis of multiple pairs of Ts21 and control lines indicate that the specification and proliferation of a specific progenitor pool (COUP-TFII+) is altered in Ts21 **(Figure 2)** and scRNAseq data identifies dysregulated WNT signaling in Ts21 progenitors (**Figure 3)**. WNT and its antagonist SHH precisely mediate the specification and proliferation of cortical interneuron progenitors (Gulacsi and Anderson, 2006; Li et al., 2009; Xu et al., 2005). Thus, the reduced proportion of COUP-TFII+ cells in Ts21 could be due to a failure to respond WNT. We therefore tested whether the expression of SHH and WNT pathway genes was reduced in Ts21 progenitors using quantitative PCR for specific pathway genes in two sets of isogenic progenitors. We confirmed that while expression of SHH pathway genes was not consistently different in Ts21 progenitors (**Figure 5A**), expression of WNT pathway genes was reduced in Ts21 progenitors **(Figure 5B).** Comparison with expression of these genes in human fetal brain indicated that the WNT target gene *AXIN2*, but not other components of the SHH or WNT pathways, was also reduced in DS fetal brain (Olmos-Serrano et al., 2016) **(Figure 5C)**. These results corroborate that Ts21 progenitors have defects in WNT signaling machinery.

**Figure 5:**
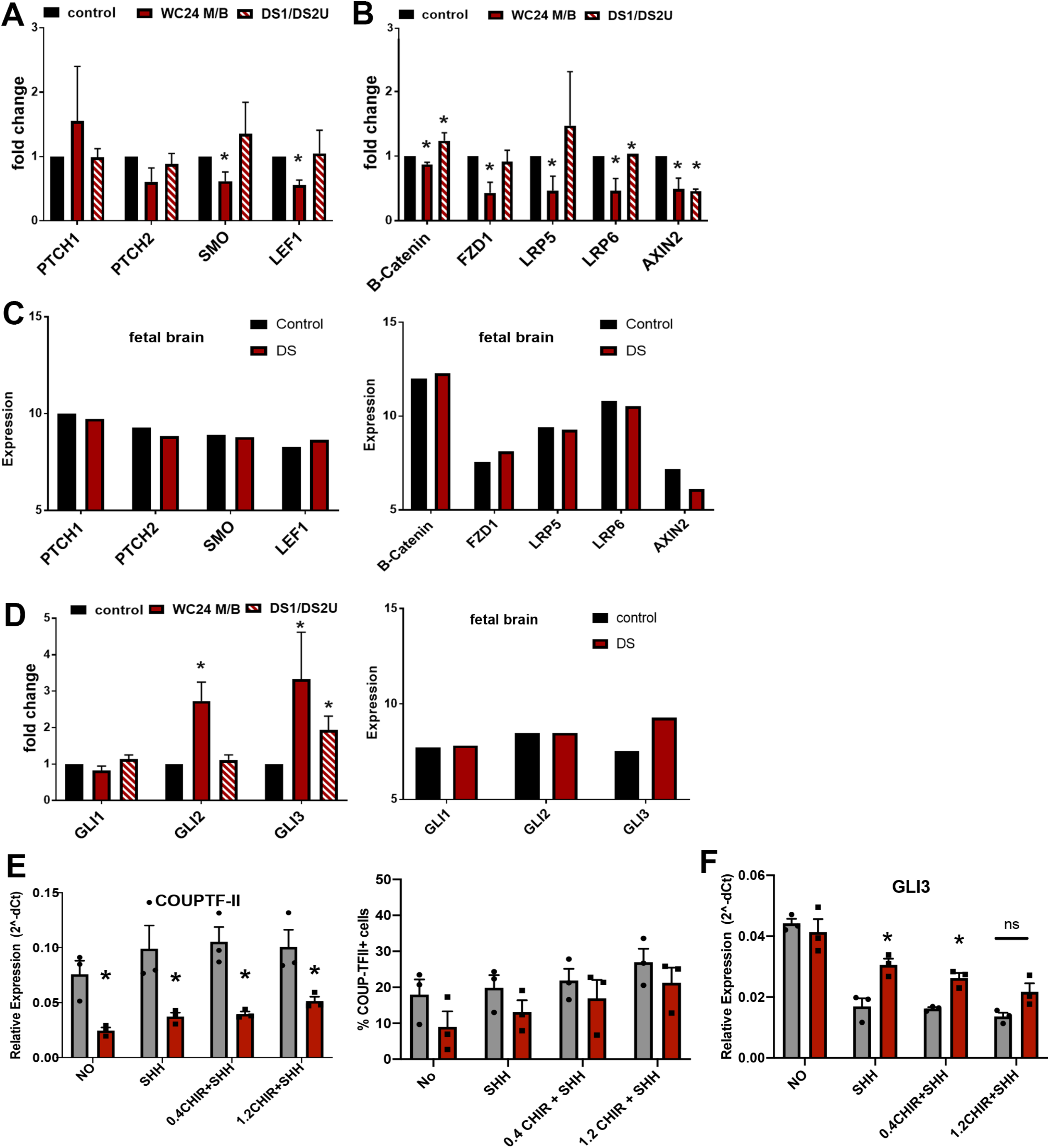
Ts21 ventral progenitors have decreased WNT and increased GLI3 expression. Quantitative PCR of SHH pathway genes (A) WNT signaling genes (B) in two iPSC lines of Ts21 and isogenic control lines. C) Expression of SHH and WNT pathway genes in human fetal control and DS brain (14-17 weeks’ gestation, dorsolateral forebrain, Olmos-Serrano et al., 2016). D) Quantitative PCR of GLI genes in progenitors cells and in human fetal control and DS brain (Olmos-Serrano et al., 2016). Statistical significance was determined by one-sample t-test on ddCt values, *p-value < 0.05, N=3 for each gene/line. E) The effects of activation of WNT via addition of a WNT agonist (CHIR, 0.4 and 1.2 um) with SHH on COUP-TFII gene expression, proportion of COUP-TFII+ cells and GLI3 expression. Statistical significance was determined by one-sample t-test on ddCt values, *p-value < 0.05, N=3 replicates in one pair of isogenic control and Ts21 cells.

We next assessed the expression of GLI genes and found that expression of *GLI3* was increased in Ts21 progenitors, in agreement with increased expression in human fetal DS brain **(Figure 5D)**. Enrichment of a *GLI3* expressing cluster of progenitors in Ts21 (**Figure 4C, E**) provides an explanation for the increased *GLI3* expression.

To test whether activation of WNT can rescue the decreased population of COUP-TFII+ cells in Ts21, we tested the response of Ts21 ventral progenitors to WNT. Activation of WNT via addition of a WNT agonist (CHIR) with SHH moderately increases both *COUP-TFII/NR2F2* expression and the proportion of COUP-TFII+ cells in Ts21 progenitors **(Figure 5E)**. Importantly, WNT activation also decreases the overexpression of *GLI3* in Ts21 progenitors (**Figure 5F**) suggesting that the enriched Cluster 3 arises from Ts21 progenitors due to decreased WNT signaling. These data support a proposed mechanism in which deficient WNT signaling in Ts21 progenitors leads to a change in specification (and possibly proliferation) of Ts21 progenitors.

## DISCUSSION

### Fewer calretinin neurons in Down syndrome

Expansion of upper layers of the cortex in primates includes more excitatory neurons in layers II and III, and there may also need to be a compensatory inhibitory neuron expansion. CR+ neurons are predominantly found in upper layers of the cortex and, in primates, a greater proportion of interneurons are CR+ neurons (Dzaja et al., 2014; Hansen et al., 2013; Hladnik et al., 2014; Ma et al., 2013). Our results align with this idea as we find a larger proportion of CR+ neurons compared to PV+ in the human STG (**Figure 1**).

By carefully assessing interneurons in post-mortem adult brain, we show that at least one interneuron subtype population, CR+, is reduced in DS. Reduced density of neurons could be due to degeneration in adulthood. However, we selected samples from 15-35 years when neuronal degeneration is not thought to take place. In fact, reduced neuron density has been reported in fetal DS brain (Golden and Hyman, 1994; Guidi et al., 2018; Larsen et al., 2008; Schmidt-Sidor et al., 1990; Stagni et al., 2018). In particular, Guidi et al., reported fewer CR+ cells in human *fetal* DS cortex in the fusiform and inferior temporal gyri (Guidi et al., 2018), corroborating our adult data and supporting the idea that this population is affected during development.

Our data are consistent with the predominant candidate mechanism underlying intellectual disability in DS that has emerged from mouse models of an imbalance in excitation-inhibition in the cortex, specifically over-inhibition, which has led to targeting of over-inhibition in the cortex by various drugs (Deidda et al., 2015; Gardiner, 2010; Martinez-Cue et al., 2014; Potier et al., 2014; Zorrilla de San Martin et al., 2018). While the decrease in interneurons that we observe may seem contrary to this hypothesis, CR+ interneurons function in disinhibitory circuits (Guet-McCreight et al., 2020; Pi et al., 2013) that specialize in inhibiting GABA interneurons. Fewer CR+ neurons in DS could provide less inhibition and thus increased activity of inhibitory GABA interneurons to elicit over-inhibition.

### Specification of interneuron progenitors is altered by Ts21

Using disorder specific iPSCs, we investigated if and how early events in interneuron development are altered in DS. At the cellular level, we corroborate that we can model the reduced generation of CR+ interneurons *in vitro*. Results indicate that NKX2.1+ MGE progenitors are not altered but fewer COUP-TFII+ progenitors are generated in Ts21. Our data appear to differ from a report that concluded that more NKX2.1+ progenitors and more interneurons are generated from Ts21 iPSCs (Xu et al., 2019). Besides difference in the use of 3D versus 2D cultures, we take an unbiased approach to assess interneuron progenitors from Ts21 iPSCs, while Xu et al., analyzed OLIG2+ cells. In addition, we are looking at an early timepoint when progenitors have established their positional identity, while Xu et al., analyzed a later timepoint. Thus, we may be assessing different progenitor populations.

In mouse CR+ interneurons derive from COUP-TFII+ progenitors and so these results link the altered development of the COUP-TFII+ subpopulation *in vitro* with fewer CR+ interneurons that we observe both i*n vitro* and *in vivo* (Huo et al., 2018). Yet, the origin of CR+ neurons has not been established in human and it is possible that some of the NKX2.1+ MGE cells also give rise to CR+ cells. CR+ neurons primarily populate upper layers of the cortex and thus likely derived from late-born COUP-TFII+ progenitors. It is also possible that specification of CR+ neurons is from MGE initially and CGE later and that decreased pools of CGE progenitors affect late born neurons.

Our data suggest potential cellular mechanisms underlying the observed decrease in COUP-TFII+ progenitors in Ts21. Ts21 progenitors could have altered specification, supported by the enrichment of a specific subpopulation of progenitors revealed by our single cell clustering. Proliferation changes during development also contribute to the decreased neuronal numbers and reduced cortical volume in DS brain (Contestabile et al., 2007; Guidi et al., 2011). The decreased number of proliferating COUP-TFII+ cells in Ts21 supports premature exit of COUP-TFII+ cells from the cell cycle. These mechanisms are not mutually exclusive, but can be addressed through lineage tracing of COUP-TFII+ progenitors.

Single-cell transcriptomics reveal that ventralized progenitors differentiated from iPSCs are heterogeneous, complicating our ability to match cellular phenotypes with molecular signatures. In particular, *COUP-TFII/NR2F2* expression in human is more complex than in mouse. Recent data from Shi et al., indicates that *PAX6, MEIS2* and *COUP-TFII/NR2F2* expression is widespread in CGE and LGE, contrary to the prevailing idea that PAX6 expression is limited to dorsal NPCs, that MEIS2 is a marker of LGE and *COUP-TFII/NR2F2* is a marker of CGE (Shi et al., 2021). Further Shi et al., uncover a cluster of *COUP-TFII/NR2F2* cells that is unique to human. A deeper understanding of the effects of Ts21 on interneuron progenitor specification requires single-cell analysis of developing DS tissue.

Cells in the Ts21-enriched cluster express markers that identify them as intermediate progenitor cells (Li et al., 2018). The presence of this cluster could indicate that Ts21 interneuron progenitors differentiate more slowly than controls and are thus retained in this intermediate stage. *GLI3* is mis-expressed temporally during fetal cortical development; *GLI3* expression is initially higher than controls and then lower than controls (Olmos-Serrano et al., 2016), consistent with altered emergence of this cluster of cells. Alternatively, this developmental cluster could represent an alternative differentiation pathway taken by Ts21 progenitors. These different scenarios can be tested through single cell analysis across developmental time, either *in vivo* or *in vitro*.

### Ts21 progenitors have reduced WNT signaling

Ts21 progenitors could have altered specification due to a reduced ability to respond to WNT, as supported by the enrichment of Cluster 3 and altered WNT pathway genes in Ts21. WNT signaling has been implicated in other cellular mechanisms associated with aging and neuropathology in DS (Adorno et al., 2018; Cairney et al., 2009; Granno et al., 2019). Since WNT is used in a tightly regulated and spatially specific manner during forebrain development to regulate regional identity, it is likely that decreased WNT signaling in Ts21 would have an impact on neurodevelopment. Our molecular data suggest that altered specification of Ts21 progenitors is at least partly due to decreased WNT signaling, which is corroborated by the restoration of the COUP-TFII+ progenitor subpopulation and *GLI3* expression through WNT activation. These data raise the hypothesis that Ts21 *GLI3* expressing intermediate progenitors in Cluster 3 emerge due to reduced WNT responsiveness. It will be important to test whether WNT activation can specifically eliminate Cluster 3 and/or whether other progenitor subpopulations are affected by WNT activation through additional single cell analyses.

## EXPERIMENTAL PROCEDURES

### Quantification of neurons in post-mortem brain

#### Tissue

Postmortem brain tissue was obtained from the NICHD Brain and Tissue Bank for Developmental Disorders (Neurobiobank) with approval from the University of Wisconsin-Madison Human Subjects IRB. Superior temporal gyrus or Brodmann’s Area 22 was obtained from 4 DS individuals and age and gender matched control subjects (**Table S1**).

#### Immunocytochemistry

Tissues were cryosectioned at 50 microns and processed for immunocytochemistry. Antigen-antibodies were visualized with avidin-biotin, horseradish peroxidase (HRP) and 3, 3’-Diaminobenzidine (DAB) using standard immunohistochemical techniques on floating sections (**Table S3**).

#### Quantification of positive cells

Total numbers of NeuN+, PV+, CR+ and SST+ neurons were estimated using the Optical Fractionator (OF) workflow in Stereo Investigator software (MBF Bioscience). The total cell population estimate (from the OF Workflow) was divided by the total tissue volume (from the Cavalieri Estimator) to calculate cell density.

### Human induced pluripotent stem cells (iPSCs)

#### iPSCs

We used two Ts21 iPSC isogenic pairs and additional iPSCs from DS individuals and unaffected controls (**Table S2**). Primary dermal fibroblasts were isolated from tissue acquired with approval from the University of Wisconsin-Madison Human Subjects IRB (protocol #2016–0979). Fibroblasts were reprogrammed by electroporation delivery of episomal vectors pCXLE-hOCT3/4-shp53-F (Addgene, 27077), pCXLE-hSK (Addgene, 27078) and pCXLE-hUL (Addgene, 27080). The iPSC colonies were manually picked between day 14-28 post-transfection. Following expansion, cells were transferred onto matrigel (R&D) and cultured with mTeSR1 (Stemcell Technologies) for banking.

#### Cell culture

iPSCs were maintained on MEFs in hESC media (DMEM/F-12/KOSR/L-Glut/MEM-NEAA/FGF-2) and passaged with collagenase. Differentiation to interneurons in vivo.ointerneuron progenitors was carried out as described (Liu et al., 2013) using SHH as a morphogen and maintaining neurospheres in NIM with B27 and purmorphamine.

### Cellular analysis

#### Cell proliferation

Cell proliferation was assayed using Click-iT™ EdU Alexa Fluor™ 488 Imaging Kit. EdU was added to cells for 8 hours. Cells were fixed with 4% paraformaldehyde in PBS for 15 minutes and processed for immunofluorescence.

#### Immunofluorescence

NPCs were plated onto laminin coated 96 well cell culture plates or coverslips at 50-60,000 cells/well/coverslip. The day after plating, cells were fixed with 4% paraformaldehyde and processed for immunofluorescence.

#### High Content Imaging analysis

Imaging and analysis was done using the high content imager Operetta (Perkin Elmer) at 20x magnification.

### Molecular analysis

#### qPCR

qPCR was performed in triplicate on 2-3 batches of differentiation (N=3) (**Table S4**). Data are presented as Fold Change calculated from ddCt values. Error bars indicate fold change of ddCt values +/-1 SD. Statistical significance was determined by one-sample t-test on ddCt values.

#### Single Cell RNA sequencing

NPCs were analyzed using the 10x Genomics Chromium Single Cell Gene Expression Assay at the University of Wisconsin Biotechnology Center. Sequence data was analyzed on servers at the UW-Madison Bioinformatics Resources Center remotely via Bash UNIX shell commands. Raw read quality was confirmed with FastQC Processing. The 10x Genomics pipeline linux commands were used for processing the data for analysis. Briefly, Cell Ranger mkfastq was used to demultiplex the raw read files in to FastQ files. The FastQ files were then passed on to Cell Ranger count for alignment, filtering, barcode counting, and UMI counting.

The processed count matrix contains 33,694 genes; and 2,134 cells in the control group and 2,158 cells in the Trisomy21 group, respectively. We used Seurat 4.0.4 for downstream analysis. First, we filtered cells that detected unique genes less than 20 or over 10^5; filtered cells that have more than 5% mitochondrial counts. After filtering, the dataset remains 4,025 cells (control: 2,003; Trisomy 21: 2,022). Second, we normalized the data and found the top 2000 highly variable genes using Seurat. We implemented integrated analysis for the two samples. Then we mitigated the effects of cell cycle heterogeneity in the dataset by calculating cell cycle phase scores based on canonical markers, and regressing these out of the data. Next, we performed PCA for dimension reduction on the integrated dataset for later clustering and visualization. Code is available in Supplemental Methods.

### Statistics

Experiments include three biological replicates (batches of differentiation, N=3 or individual cell lines N=3 or 4) and 3 technical replicates (n=3) for each cell line. Ts21 and control pairs were differentiated together. Data were analyzed using GraphPad Prism version 8. All pooled data are presented as mean ± standard error of the mean (SEM). Differences were considered statistically significant at p<0.05.

## ACKNOWLEDGEMENTS

We thank members of the Bhattacharyya and Zhang labs for helpful comments and technical assistance. We thank Karla Knobel, Emily Fares, Anna Baker, MBF technical support, Manuel Casanova and Kenneth Fish for guidance on stereology and S. Splinter-BonDurant, D. Pavelec, and M.E. Berres at the UW-Madison Biotechnology Center for technical assistance. This work was supported by NIH grants R03HD083538 and R21NS105339 to A.B., 1R01HD106197 to A.B and S-C.Z., and funding from UW-Madison and the Wisconsin Alumni Research Foundation to A.B. and in part by a core grant to the Waisman Center from the National Institute of Child Health and Human Development (U54 HD090256). Zhang is co-founder of BrainXell, Inc. The authors declare no competing interests.

## AUTHOR CONTRIBUTIONS

Conceptualization, A.B. and S-C.Z.; Methodology, Y.G-R., J.S., B.S., M.M. D.W.; Software and Data Curation, Y.G-R., J.S., D.W.; Investigation, Y.G-R., J.S. B.S., K.X., L.H., M.M., K.A.R., S.D., S.M., A.K., R.A.D., B.L., L.A., R.R.; Writing – Original Draft, Y.G-R., A.B.; Writing – Review & Editing, A.B., Y.T., A.S., S-C.Z.; Funding Acquisition, Project Administration, Resources and Supervision, A.B., S-C.Z., D.W.

