## supplemental for "Altered patterning of trisomy 21 interneuron progenitors"

Figure S1, related to Figure 3

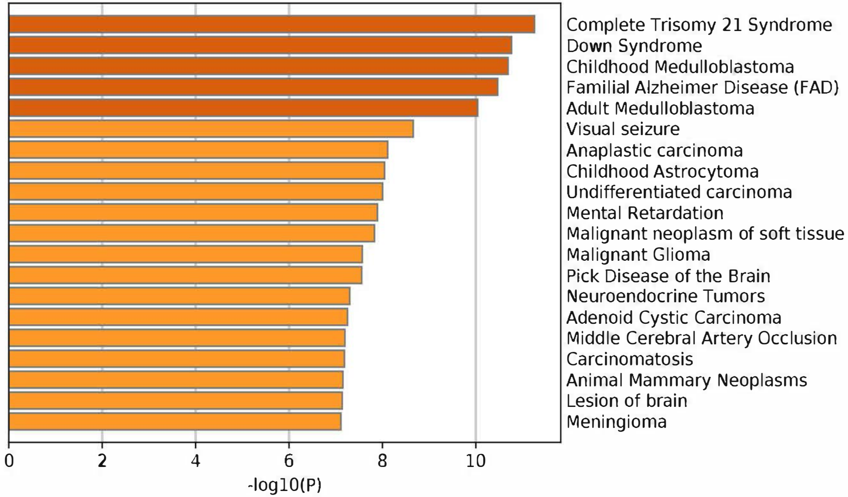


DEG pathways identified by DisGeNet database.

### Figure S2, related to Figure 3

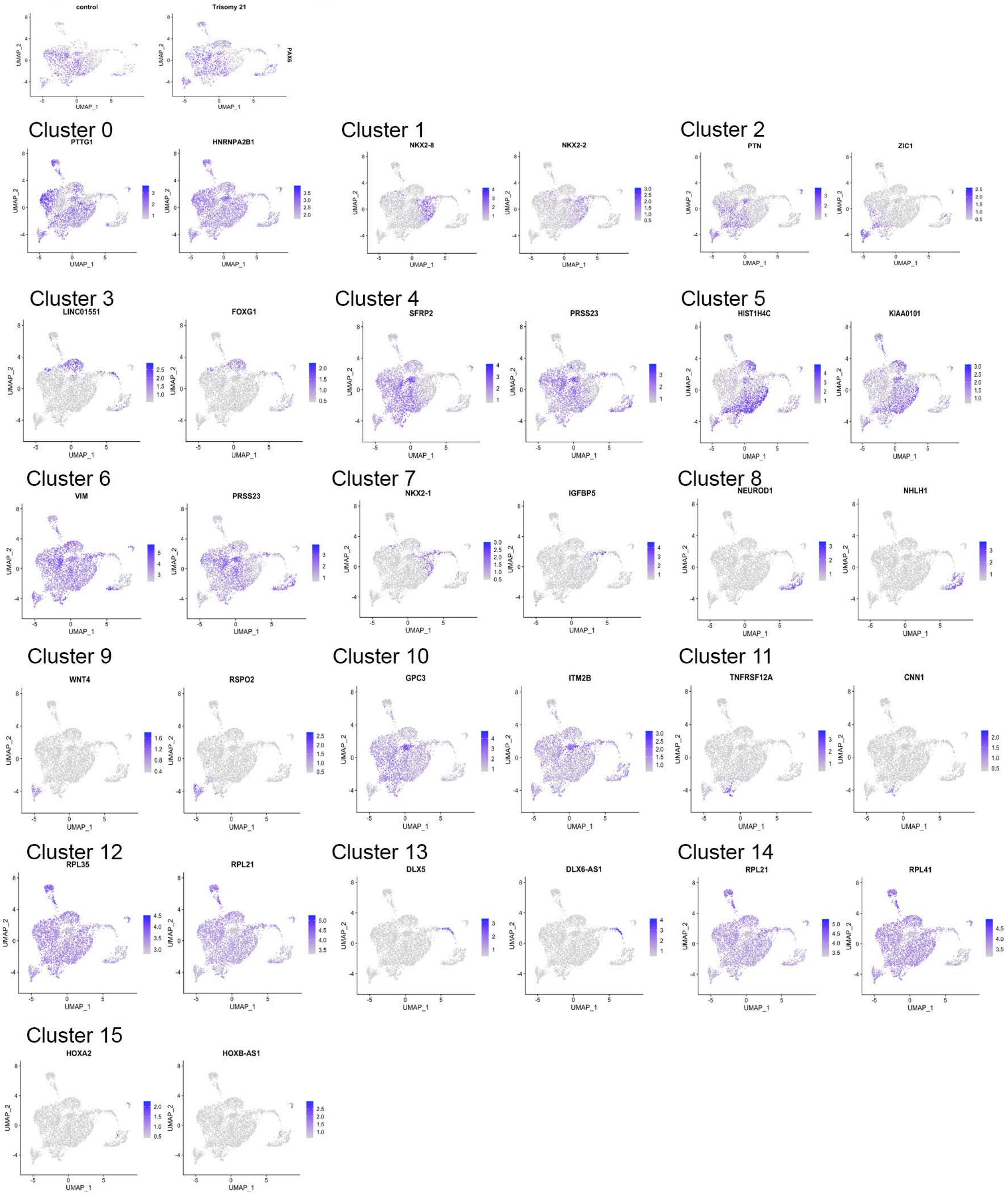

Figure S3, related to Figure 4

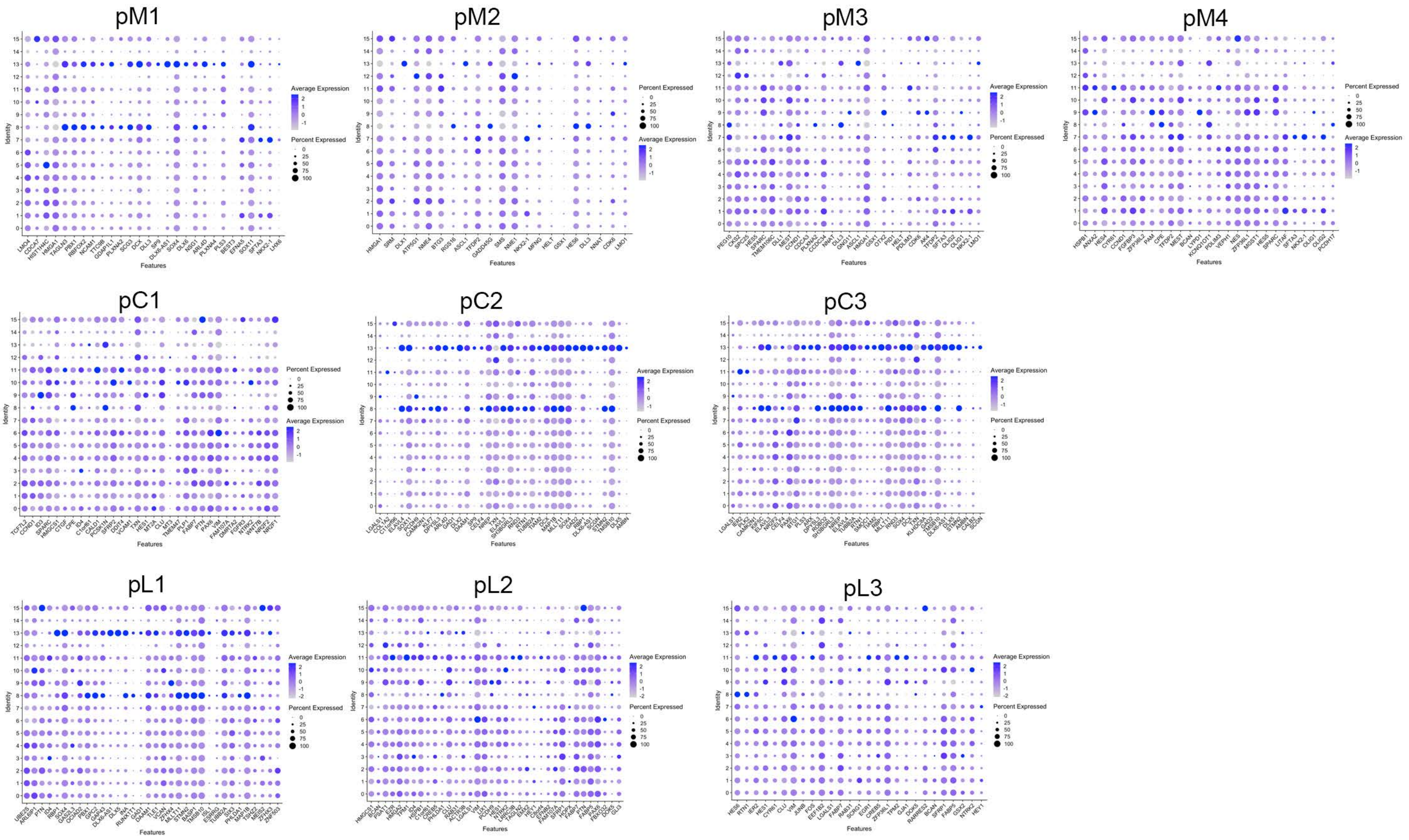


Dot plots comparing gene expression in iPSCs clusters (Figure 4) (y-axis) to gene expression in fetal ganglionic eminences subclusters (Shi et al., 2021) (x-axis).

**Table S1: Subject and sample information**

| UMB number | Diagnosis | Age (years, days) | Sex | Race | Post mortem interval (hours) |
|---|---|---|---|---|---|
| 1841 | Control | 19, 289 | Male | Caucasian | 14 |
| 5277 | Ts21 | 19, 352 | Male | Caucasian | 26 |
| 5654 | Control | 19, 264 | Male | Caucasian | 18 |
| M1960M | Ts21 | 19, 311 | Male | Caucasian | 14 |
| 5030 | Control | 24, 333 | Male | Afr Amer | 14 |
| 5341 | Ts21 | 25, 304 | Male | Afr Amer | 24 |
| 1544 | Control | 32, 315 | Male | Caucasian | 12 |
| 4273 | Ts21 | 33, 315 | Male | Caucasian | 36 |

**Table S2: iPSCs used in this study**

<table>
<tr><th>Cell line</th><th>Sex</th><th>Age (years)</th><th>Karyotype</th><th>Relationship</th><th>Reprogramming method</th></tr>
<tr><td>WC-24-B</td><td rowspan="2">Female</td><td rowspan="2">25</td><td>Normal</td><td rowspan="2">Isogenic pair</td><td>Episomal</td></tr>
<tr><td>WC-24-M</td><td>Trisomy 21</td><td></td></tr>
<tr><td>DS2U</td><td rowspan="2">Male</td><td rowspan="2">1</td><td>Normal</td><td rowspan="2">Isogenic pair</td><td>Retroviral</td></tr>
<tr><td>DS1</td><td>Trisomy 21</td><td></td></tr>
<tr><td>603-8</td><td>Male</td><td>36</td><td>Normal</td><td rowspan="2">Unrelated</td><td>Retroviral</td></tr>
<tr><td>WC-38-01</td><td>Male</td><td>35</td><td>Trisomy 21</td><td>Sendai</td></tr>
<tr><td>WC-58-07</td><td>Female</td><td>Neonate</td><td>Normal</td><td rowspan="2">Unrelated</td><td>Episomal</td></tr>
<tr><td>WC-20-02</td><td>Female</td><td>3</td><td>Trisomy 21</td><td>Episomal</td></tr>
</table>

**Table S3: Immunocytochemical methods**

| MARKER | | ANTIBODY INFORMATION | ANTIGEN RETRIEVAL | DILUTION | secondary, ABC | DAB |
|---|---|---|---|---|---|---|
| all neurons | NeuN | AbCam ab104225 Rabbit | Vector unmasking 15 minutes 95°C | 1:500 | Visucyte HRP polymer | 5 minutes ImPACT |
| Parvalbumin | PV | Sigma P3088 mouse | Vector unmasking 5 minutes 95°C | 1:1000 | Biotin 2' ABC | 5 minutes |
| Somatostatin | SST | Millipore MAB354 rat | Vector unmasking 15 minutes 95°C | 1:100 | Biotin 2' ABC | 10 minutes |
| Calretinin | CR | Swant CR7697 Rabbit | Vector unmasking 15 minutes 95°C | 1:2000 | Biotin 2' ABC | 3 minutes |

**Table S4: Antibodies used in immunofluorescence.**

| Antigen | Company and catalog number |
|---|---|
| NKX2.1/TTF1 | abCAM ab76013 |
| COUP-TFII/NRF2 | R&D systems PP-H7147-00 |
| Calretinin | Swant 7697 |
| Somatostatin | Millipore MAB354 |
| Phospho-Histone H3 (Ser10) | Cell Signaling Technology 9706 |

**Table S5: Primer list**

| Gene | Forward Seq (5' – 3') | Reverse Seq (5' – 3') |
|---|---|---|
| AXIN2 | TATCCAGTGATGCGCTGA | CGGTGGGTTCTCGGGAAATG |
| NR2F2/COUPTFII | CTCAAGGCCATAGTCCTGTCC | GGTACTGGCTCCTAACGTATTC |
| FZD1 | ATCTTCTTGTCCGGCTGTTACA | GTCCTCGGCGAACTTGTCATT |
| GLI1 | AACGCTATACAGATCCTAGCTCG | GTGCCGTTTGGTCACATGG |
| GLI2 | CCCCTACCGATTGACATGCG | GAAAGCCGGATCAAGGAGATG |
| GLI3 | GAAGTGCTCCACTCGAACAGA | GTGGCTGCATAGTGATTGCG |
| LEF1 | ATGTCAACTCCAAACAAGGCA | CCCGGAGACAAGGGATAAAAAGT |
| LRP5 | CGACACTGGGACCAACAGAA | AGATGTAGCCCTTGGTGGGA |
| LRP6 | CTGAGAGCGGCCCCTTTGTT | GCATCCTCCAAGCCTCCAAC |
| NKX 2.1 | AGCACACGACTCCGTTCTC | GCCCACTTTCTTGTAGCTTTCC |
| PTCH1 | GGAGCAGATTTCCAAGGGGA | CCACAACCAAGAACTTGCCG |
| PTCH2 | CCGCCAGAGGTGATACAGAT | CCACGGTCATGGAGGTAGTC |
| SMO | ACTTGGATTGCGAGGCTAGG | TCGCAAACTTTGGAACCCG |

**Supplemental Experimental Procedures**

**Quantification of neurons in post-mortem brain**

*Tissue:* Adult postmortem brain tissue was obtained from the NICHD Brain and Tissue Bank for Developmental Disorders with approval from the University of Wisconsin-Madison Institutional Review Board (**Table S1**). Superior temporal gyrus or Brodmann's Area 22 was obtained from four DS individuals and age and gender matched control subjects. It should be noted that the post mortem interval (PMI) varied between the samples and, in particular, the DS samples had longer PMIs than their matched controls (Controls 14.50 $\pm$ 1.26, DS 25.00 $\pm$ 4.5, p=0.11 calculated using non-parametric Mann-Whitney U test).

*Immunocytochemistry:* Tissues were sectioned at 50 microns using a cryostat and processed for immunocytochemistry. Antigen-antibodies were visualized with avidin-biotin, horseradish peroxidase (HRP) and 3, 3'-Diaminobenzidine (DAB) using standard immunohistochemical techniques on floating sections (**Table S3**).

*Quantification of positive cells:* Total numbers of NeuN+, PV+, CR+ and SST+ neurons were estimated using the Optical Fractionator (OF) workflow in Stereo Investigator software (MBF Bioscience). Six to eight sections at an interval of 5-7 were analyzed. Percentages of tissue for sampling were chosen based on the resample oversample function in Stereo Investigator (PV: 1%, CR: 1%) to ensure a coefficient of error < 0.1.The total positive cell count was estimated using the following equation: $N = \sum Q * \frac{t}{h} * \frac{1}{asf} * \frac{1}{ssf}$, where $\sum Q$is the total number of cells counted, *t* the average section thickness and *h* the height of the optical dissector, and *asf* and *ssf* the area sectioning fraction and the section sampling fractions, respectively [10].

*Volume and density calculation:* Total volume of tissue sampled and counted was estimated using the Cavalieri Estimator probe (Stereo Investigator, MBF Bioscience). The total cell population estimate (from the Optical Fractionator Workflow) was divided by the total tissue volume (from the Cavalieri Estimator) to calculate cell density.

**Human induced pluripotent stem cells (iPSCs)**

*iPSCs:* We established one new isogenic Ts21 iPSC pair and additional iPSCs from DS individuals and unaffected controls (**Table S2**). Primary dermal fibroblasts were isolated from tissue acquired with approval from the University of Wisconsin-Madison Human Subjects Institutional Review Board (protocol #2016–0979). Fibroblasts were reprogrammed by electroporation delivery of episomal vectors pCXLE-hOCT3/4-shp53-F (Addgene, 27077), pCXLE-hSK (Addgene, 27078) and pCXLE-hUL (Addgene, 27080). After electroporation, cells were cultured on mouse embryonic fibroblast (MEF) feeder cells in a low oxygen incubator (5% $O_2$, 5% $CO_2$). Cells were fed with hESCM (DMEM-F12 media (Gibco) with 20% knock-out serum replacement (Gibco), 1X Non-Essential Amino Acids (Life Technologies), 0.5X GlutaMAX (Life Technologies), 0.1 mM 2-mercaptoethanol (Sigma), and 12 ng/mL bFGF (Waisman Biomanufacturing)). The iPSC colonies were manually picked between day 14–28 post-transfection. Following expansion, cells were transferred onto Matrigel (R&D) and cultured with mTeSR1 (Stemcell Technologies) for banking. iPSCs on MEF were passaged with dispase solution (Gibco) Split ratio is 1 to 6 every 5–7 days., and iPSCs on Matrigel were passaged with 0.5mM EDTA or ReLeSR (05872, Stemcell Technologies).

*Cell culture:* iPSCs were maintained on MEFs in hESC media (DMEM/F-12/KOSR/L-Glut/MEM-NEAA/FGF-2) and passaged with collagenase. Differentiation to interneuron progenitors was carried out as described (Liu et al., 2013). When ~80% confluent, iPSCs were dissociated from MEFs using dispase to generate embryoid bodies (EBs). EBs were maintained in suspension with dual SMAD inhibition for 4 days and then media was changed to neural induction media (NIM; DME/F12 media with N2, NEAA and heparin). EBs were plated on Day 7 and SHH was added on Day 10. For neurons, EBs were lifted to neurospheres on Day 15 or 16 and maintained in NIM with B27 and purmorphamine. For neuronal differentiation, progenitors in neurospheres were dissociated with Accutase and plated on polyornithine/laminin-coated coverslips in neural differentiation medium containing DMEM/F12, N2(1:50), B27 (1:100), 10 ng/mL brain derived neurotrophic factor (BDNF) (Peprotech), 10 ng/mL glial derived neurotrophic factor(GDNF) (R&D Systems), cAMP (Sigma), and ascorbic acid(Sigma). Compound E (gamma secretase inhibitor XXI) was added at plating.

**Cellular analysis**

*Cell proliferation:* Cell proliferation was assayed using Click-iT™ EdU Alexa Fluor™ 488 Imaging Kit. A final concentration of 10uM EdU was added to cells for 8 hours. Cells were fixed with 4% paraformaldehyde in PBS for 15 minutes and processed for immunofluorescence for phospho-histone H3 (pHH3, Cell Signaling Technology #9706).

*Immunofluorescence:* Neural progenitors in neurospheres were dissociated with Accutase and plated onto laminin coated 96 well cell culture plates or coverslips at 50-60,000 cells/well/coverslip. The day after plating, cells were fixed with 4% paraformaldehyde in PBS for 15 minutes. Cells were rinsed with PBS and incubated with a permeabilization/blocking buffer (5% normal goat serum, 0.1% TritonX-100 in PBS) for 30 minutes. Cells were incubated with primary antibodies to NKX2.1 and/or COUP-TFII (**Table S4**) overnight followed by fluorescent secondary antibodies, washed with PBS and mounted in Fluoromount.

*High Content Imaging analysis:* Imaging and analysis was done using the high content imager Operetta (Perkin Elmer) at 20x magnification.

**Molecular analysis**

*qPCR:* RNA was isolated from progenitors using the ZYMO Research Direct-Zol RNA Miniprep plus kit followed by cDNA synthesis using qScript cDNA Supermix. qPCR was done in triplicate on 2-3 batches of differentiation (N=3) (**Table S5**). Data are presented as Fold Change calculated from ddCt values. Error bars indicate fold change of ddCt values +/- 1 SD. Statistical significance was determined by one-sample t-test on ddCt values.

*Single Cell RNA sequencing analysis:*

WC24BM_02062022.html

**Quantification and Statistical Analysis**

All experiments include at least three biological replicates (batches of differentiation, N=3 or individual cell lines N=4) and 3 technical replicates (n=3) for each cell line. Ts21 and control pairs were differentiated together. Data were analyzed using GraphPad Prism version 8. All pooled data are presented as mean ± standard error of the mean (SEM). Details regarding number of technical and biological replicates are provided in the figure legends with specific statistical analysis test used. For parametric datasets, data were analyzed using unpaired two-tailed Student's t-test. For non-parametric datasets, an unpaired Mann-Whitney test was performed. ANOVA analyses were used for datasets with more than two groups. Kruskal-Wallis analysis of variance, one-way ANOVA followed by Dunn's post hoc or Dunnett's post hoc analysis or two-way ANOVA followed by post hoc Sidak's test or Tukey's test (GraphPad Prism 8). Differences were considered statistically significant at $p<0.05$.

### R Notebook

### 1 Prepare dataset

Load dataset. There are $4,292$ single cells and $33,694$ genes. The dataset contains two experimental groups:

- Control: 2,134 cells.
- Trisomy 21: 2158 cells.

```
library(dplyr)
library(Seurat)
library(patchwork)

rm(list=ls())
setwd("~/Desktop/Waisman/Anita/filtered_gene_bc_matrices_mex/GRCh38/")
mat = read.csv("./data.csv",header = FALSE)
features = read.table("./genes.tsv",sep='\t')
barcodes = read.table("./barcodes.tsv",sep='\t')
group = read.csv('./group.csv',row.names = 1)
colnames(mat) = features$V2
rownames(mat) = barcodes$V1

seuobj = CreateSeuratObject(t(mat),min.cells = 0)
seuobj <- AddMetaData(object = seuobj, metadata = group, col.name = 'group')
```

### 2 QC and selecting cells for further analysis

The three QC metrics shown in the following violin figures are:

- The number of unique genes detected in each cell:
    - Low-quality cells or empty droplets will often have very few genes;
    - Cell doublets or multiplets may exhibit an aberrantly high gene count.
- The total number of molecules detected within a cell.
- The percentage of reads that map to the mitochondrial genome:
    - Low-quality / dying cells often exhibit extensive mitochondrial contamination.

```
seuobj[["percent.mt"]] <- PercentageFeatureSet(seuobj, pattern = "^MT-")
VlnPlot(seuobj, features = c("nFeature_RNA", "nCount_RNA", "percent.mt"), ncol = 3)
```

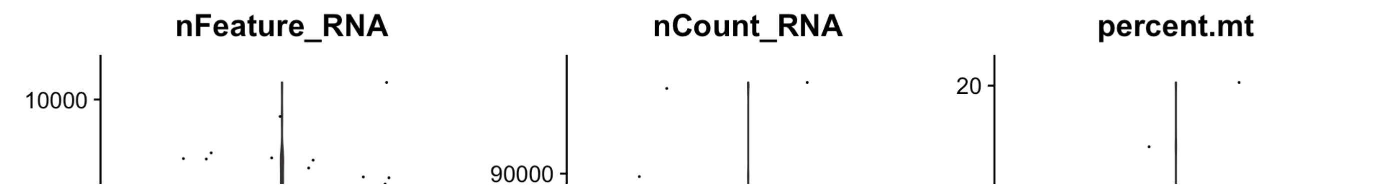

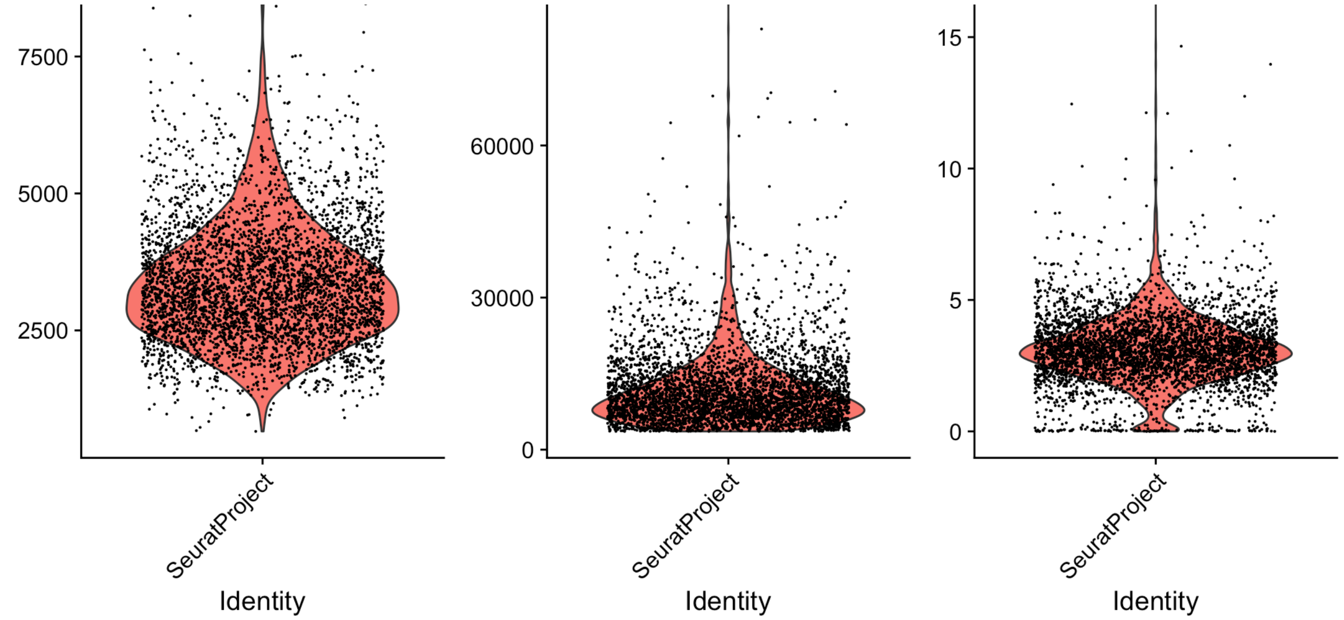


Filter cells that have detected unique genes less than $20$ or over $10^5$; filter cells that have $> 5\%$ mitochondrial counts. After filtering, the dataset remains $4,025$ cells (control: $2,003$; Trisomy 21: $2,022$).

```
seuobj <- subset(seuobj, subset = nFeature_RNA > 20 & nFeature_RNA < 10000 & percent.
mt < 5)
```

### 3 Perform integration

The joint analysis of two or more single-cell datasets might be problematic, especially identifying cell populations. We implement integrated analysis across different datasets to correct for technical differences between datasets (i.e., batch effect correction) and perform comparative scRNA-seq analysis across experimental conditions. First, we show the UMAP of all cells before integration, colored by experimental groups:

### UMAP (Dataset before integrating)

```
seuobj <- NormalizeData(seuobj, verbose = FALSE)
seuobj <- FindVariableFeatures(seuobj, selection.method = "vst", nfeatures = 2000, ve
rbose = FALSE)
seuobj <- ScaleData(seuobj, verbose = FALSE)
seuobj <- RunPCA(seuobj, features = VariableFeatures(seuobj), verbose = FALSE)
seuobj <- RunUMAP(seuobj, reduction = "pca", dims = 1:30, verbose=FALSE)
DimPlot(seuobj, group.by = "group")
```

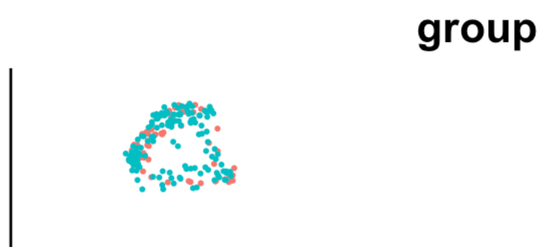

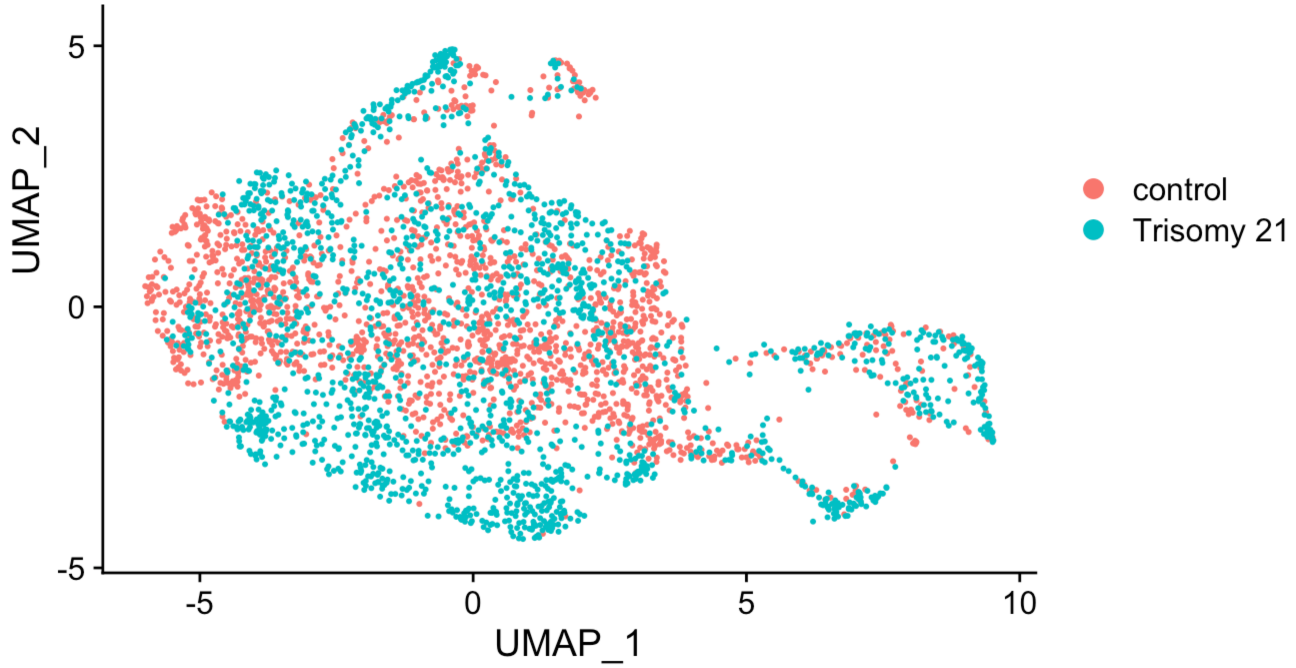


### Integration

```
seuobj.list <- SplitObject(seuobj, split.by = "group")
seuobj.list <- lapply(X = seuobj.list, FUN = function(x) {
  x <- NormalizeData(x, verbose = FALSE)
  x <- FindVariableFeatures(x, selection.method = "vst", nfeatures = 2000, verbose =
FALSE)
})
features <- SelectIntegrationFeatures(object.list = seuobj.list)
seuobj.list <- lapply(X = seuobj.list, FUN = function(x) {
  x <- ScaleData(x, features = features, verbose = FALSE)
  x <- RunPCA(x, features = features, verbose = FALSE)
})
anchors <- FindIntegrationAnchors(object.list = seuobj.list, anchor.features = featur
es, verbose = FALSE)
combined <- IntegrateData(anchorset = anchors, verbose = FALSE)
DefaultAssay(combined) <- "integrated"
combined <- ScaleData(combined, verbose = FALSE)
```

Then, the following plot shows the UMAP of all cells after integration, colored by experimental groups:

### UMAP (Dataset after integrating)

```
combined <- RunPCA(combined, features = VariableFeatures(combined),verbose = FALSE)
combined <- RunUMAP(combined, reduction = "pca", dims = 1:30, verbose=FALSE)
DimPlot(combined, group.by = "group")
```

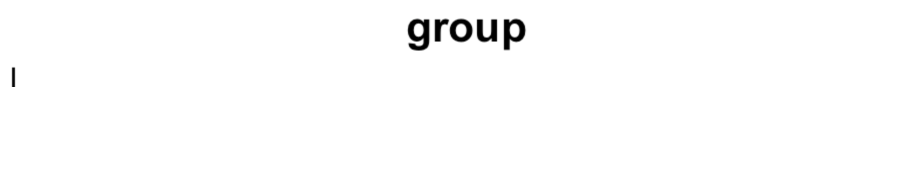

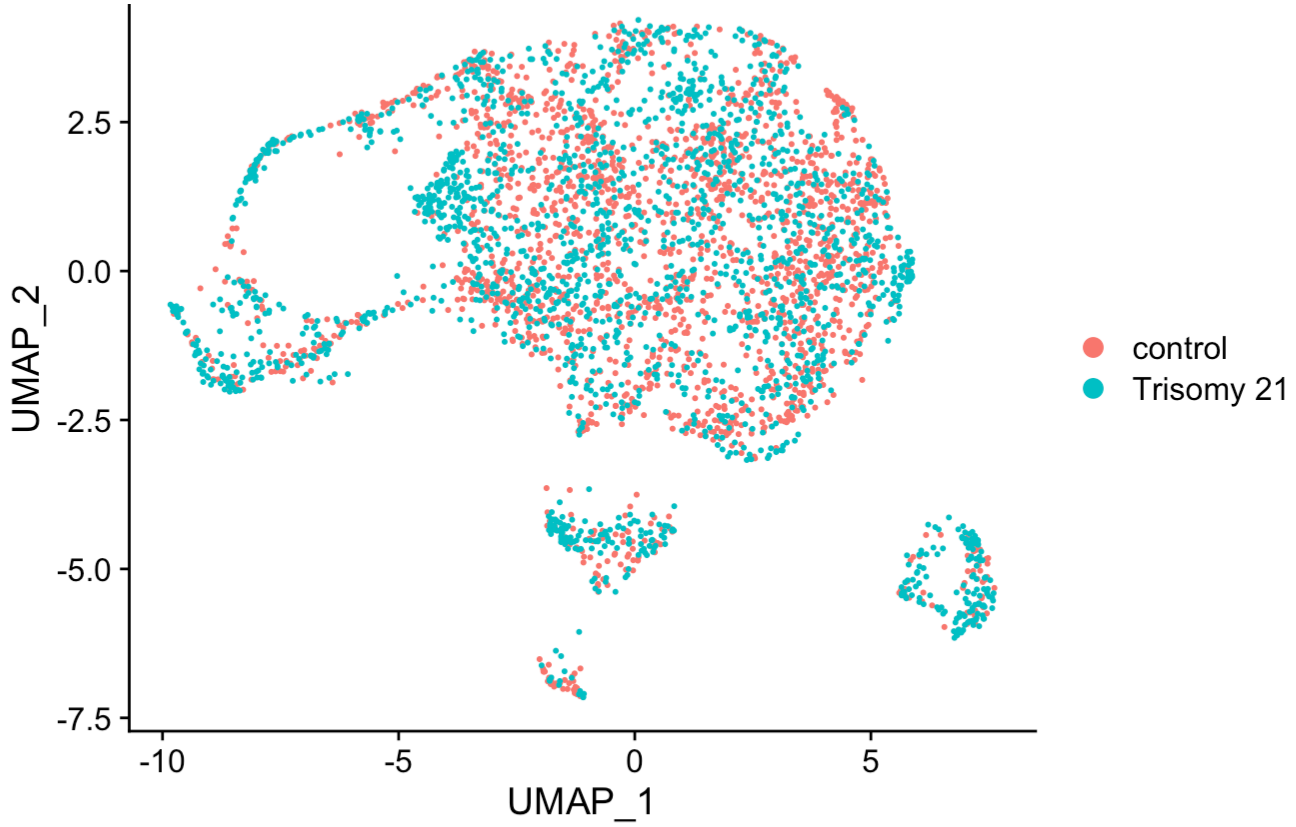


### 4 Regress out cell cycle genes

Then we mitigate the effects of cell cycle heterogeneity in the dataset by calculating cell cycle phase scores based on canonical markers, and regressing these out of the data during pre-processing.

### Before cell cycle

Running a PCA on cell cycle genes reveals that cells separate entirely by phase.

```
s.genes <- cc.genes$s.genes
g2m.genes <- cc.genes$g2m.genes
combined <- CellCycleScoring(combined, s.features = s.genes, g2m.features = g2m.gene
s, set.ident = TRUE, verbose = FALSE)
combined <- RunPCA(combined, features = c(s.genes, g2m.genes), verbose = FALSE)
p1=DimPlot(combined,reduction="pca")
p2=DimPlot(combined,reduction="pca",group.by = "group")
p1+p2
```

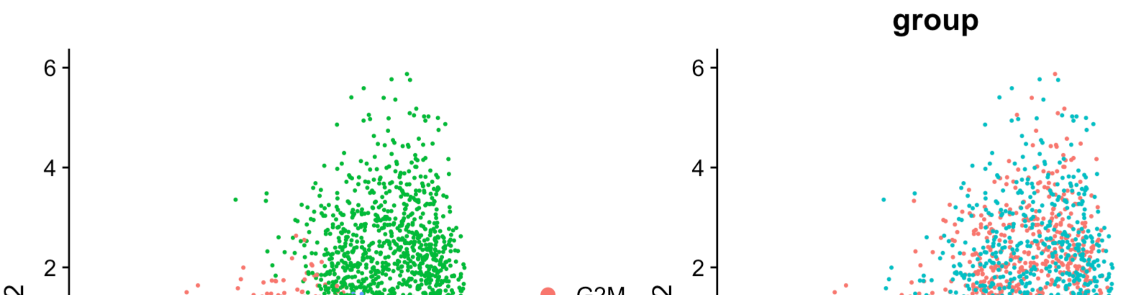

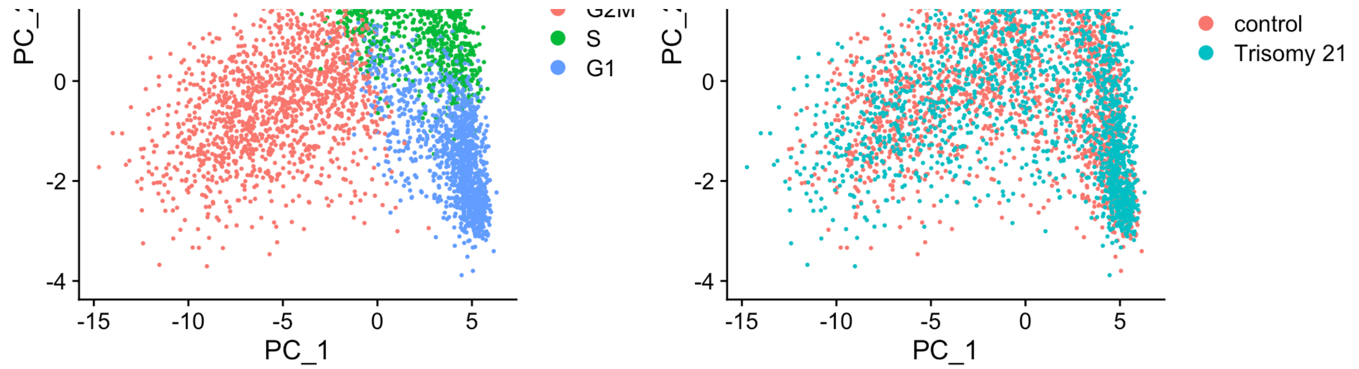


### After cell cyle

When running a PCA on only cell cycle genes, cells no longer separate by cell-cycle phase.

```
combined <- ScaleData(combined, vars.to.regress = c("S.Score", "G2M.Score"), features
= rownames(combined), verbose=FALSE)
combined <- RunPCA(combined, features = c(s.genes, g2m.genes),verbose = FALSE)
DimPlot(combined, reduction="pca")
```

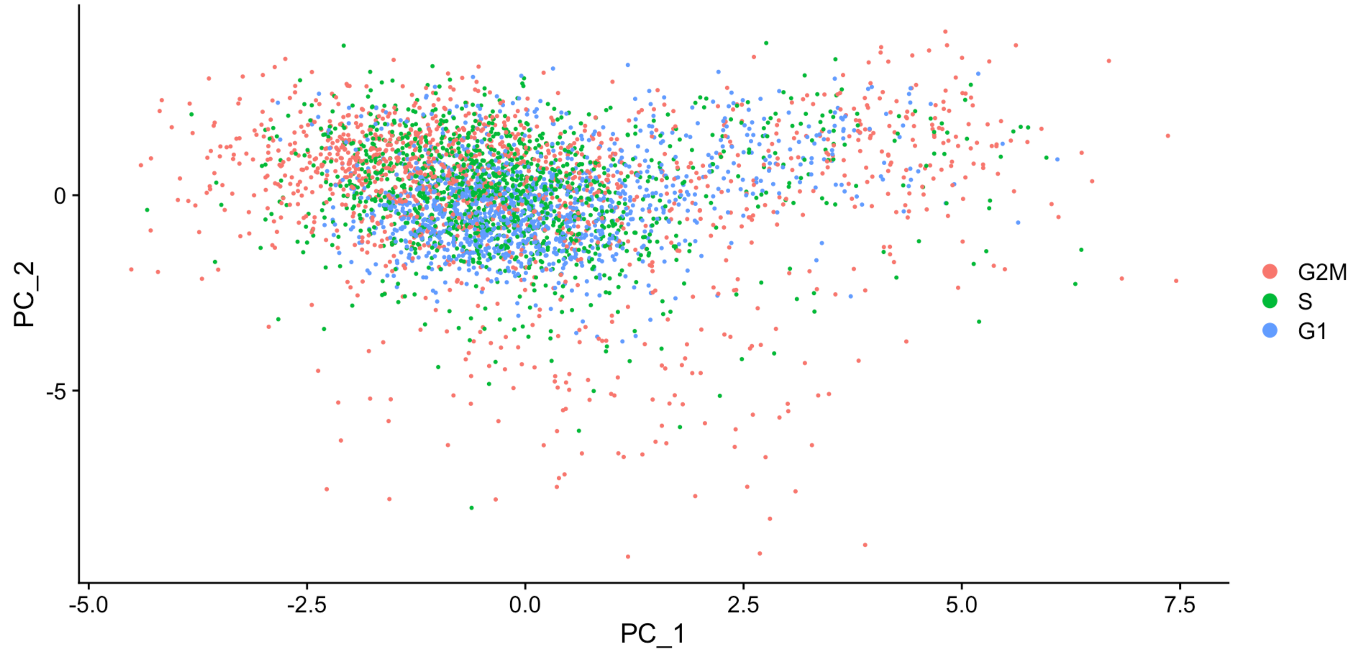


```
p1=DimPlot(combined,reduction="pca")
p2=DimPlot(combined,reduction="pca",group.by = "group")
p1+p2
```

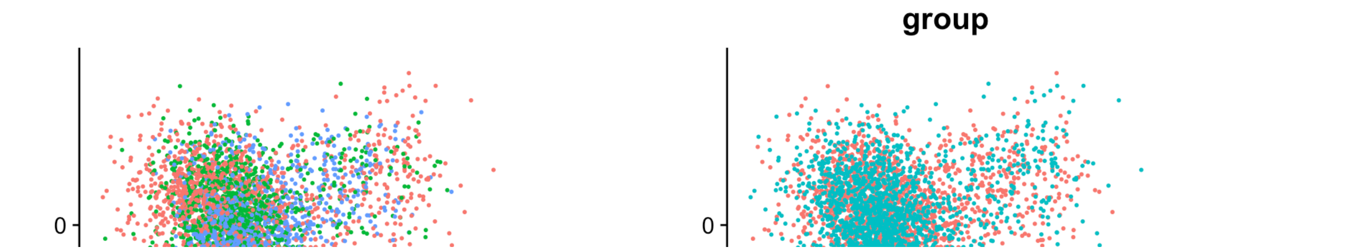

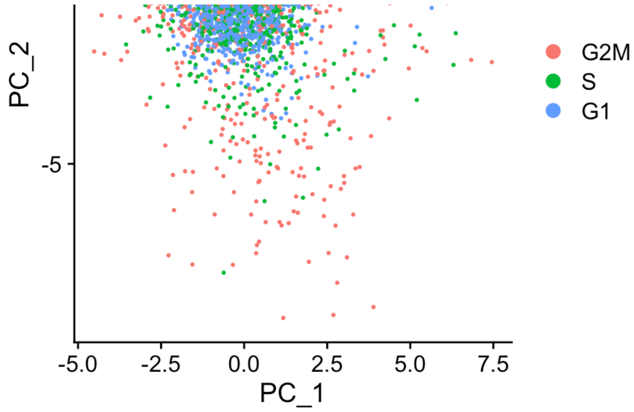


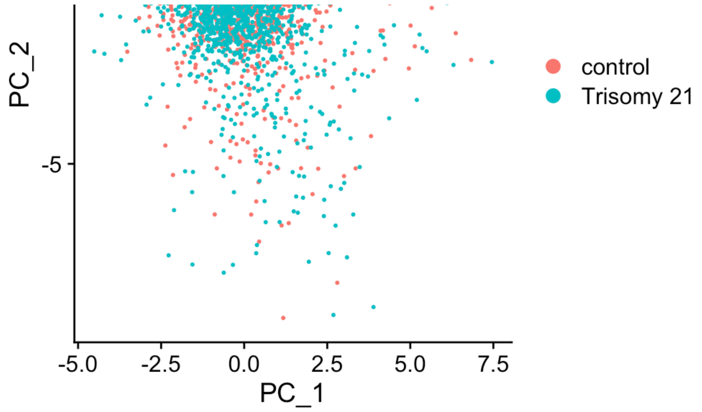


### 5 Dimensional reduction and clustering

Next, we perform PCA on the scaled data. By default, only the previously determined variable features are used as input. Then we visualize the dataset using UMAP after integration and mitigating cell cycle effects:

```
combined <- RunPCA(combined, features = VariableFeatures(combined), verbose = FALSE)
combined <- RunUMAP(combined, reduction = "pca", dims = 1:50, verbose = FALSE)
DimPlot(combined,pt.size = 0.3, group.by = "group")
```

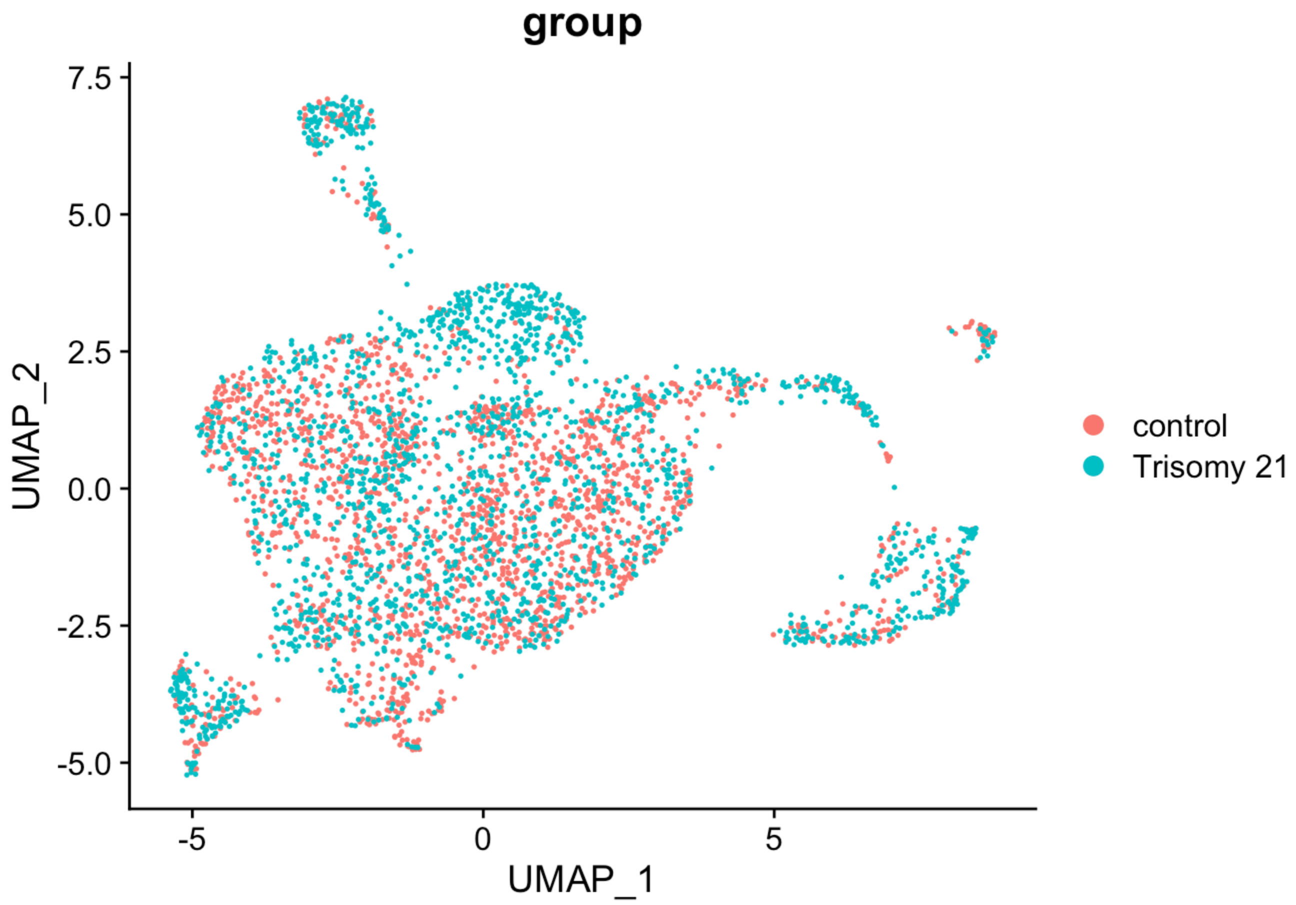


#### 5.1 Cluster the cells

Visualize by experimental groups (left) and clustering results (right):

```
combined <- FindNeighbors(combined, reduction = "pca", dims = 1:50, verbose = FALSE)
combined <- FindClusters(combined, resolution = 1, verbose = FALSE)
p1 <- DimPlot(combined, pt.size = 0.3, reduction = "umap", group.by = "group")
p2 <- DimPlot(combined, pt.size = 0.3, reduction = "umap", label = TRUE, repel = TRU
E)
p1 + p2
```

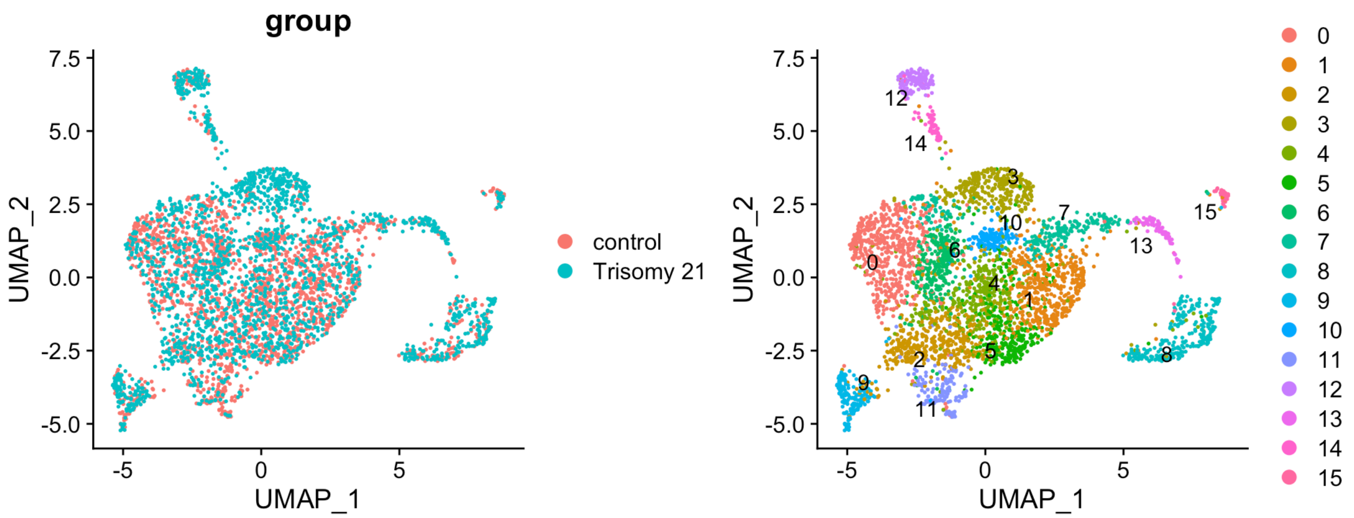


Visualize the two experimental conditions side-by-side (left: control; right: Trisomy 21):

```
DimPlot(combined, reduction = "umap", split.by = "group", label = TRUE, repel = TRUE)
```

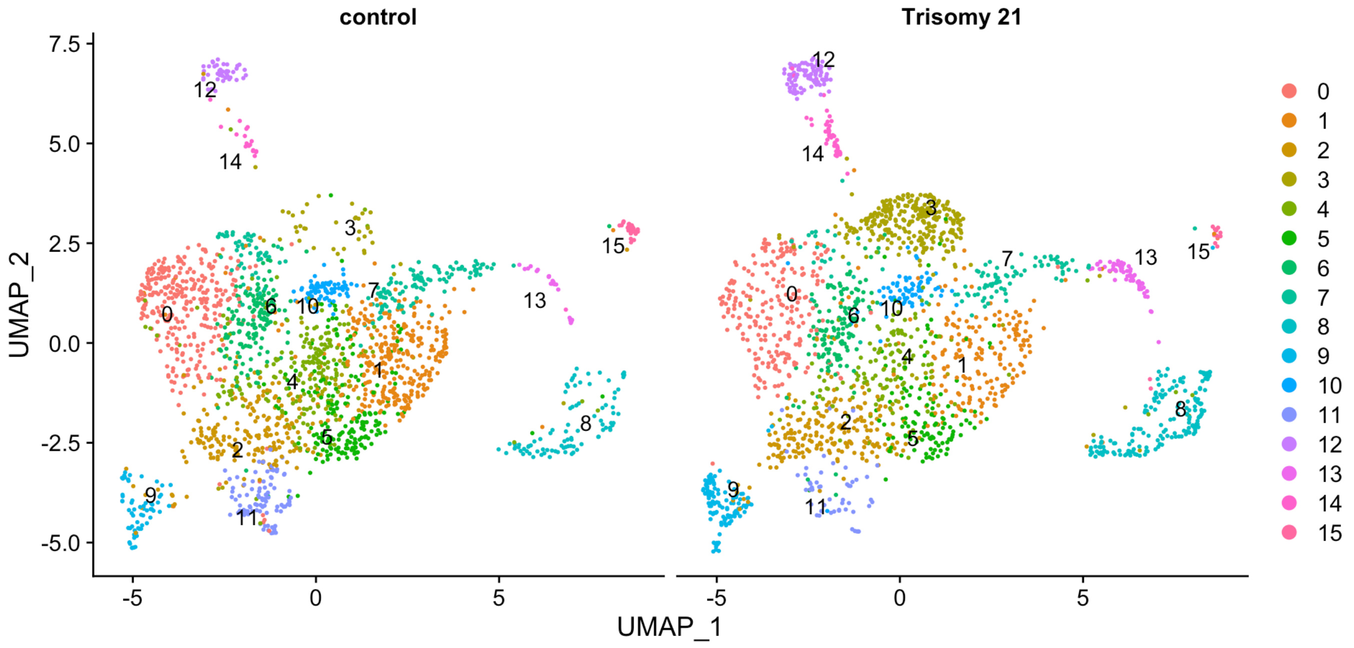


Check the number of cells in each cluster:

```
ncontrol=table(combined@meta.data[["seurat_clusters"]],combined@meta.data[["grou
p"]])[,1]
nTrisomy21=table(combined@meta.data[["seurat_clusters"]],combined@meta.data[["grou
p"]])[,2]
data.frame(`control`=ncontrol, `Trisomy 21`=nTrisomy21)
```

| | control<br><int> | Trisomy.21<br><int> |
|---|---|---|
| 0 | 289 | 223 |
| 1 | 312 | 187 |
| 2 | 208 | 210 |
| 3 | 54 | 289 |
| 4 | 201 | 130 |
| 5 | 194 | 120 |
| 6 | 157 | 139 |
| 7 | 163 | 106 |
| 8 | 90 | 175 |
| 9 | 60 | 115 |

1-10 of 16 rows Previous **1** 2 Next

Check the percentage of cells in each cluster:

```
ncontrol=table(combined@meta.data[["seurat_clusters"]],combined@meta.data[["grou
p"]])[,1]
nTrisomy21=table(combined@meta.data[["seurat_clusters"]],combined@meta.data[["grou
p"]])[,2]
data.frame(`control (%)`=ncontrol/(ncontrol+nTrisomy21)*100, `Trisomy 21 (%)`=nTrisom
y21/(ncontrol+nTrisomy21)*100)
```

| | control....<br><dbl> | Trisomy.21....<br><dbl> |
|---|---|---|
| 0 | 56.44531 | 43.55469 |
| 1 | 62.52505 | 37.47495 |
| 2 | 49.76077 | 50.23923 |
| 3 | 15.74344 | 84.25656 |
| 4 | 60.72508 | 39.27492 |
| 5 | 61.78344 | 38.21656 |

| | control.... | Trisomy.21.... |
|---|---|---|
| | <dbl> | <dbl> |
| 6 | 53.04054 | 46.95946 |
| 7 | 60.59480 | 39.40520 |
| 8 | 33.96226 | 66.03774 |
| 9 | 34.28571 | 65.71429 |

1-10 of 16 rows Previous **1** 2 Next

```
barplot(t(table(combined@meta.data[["seurat_clusters"]],combined@meta.data[["grou
p"]])), beside = TRUE, legend = TRUE)
```

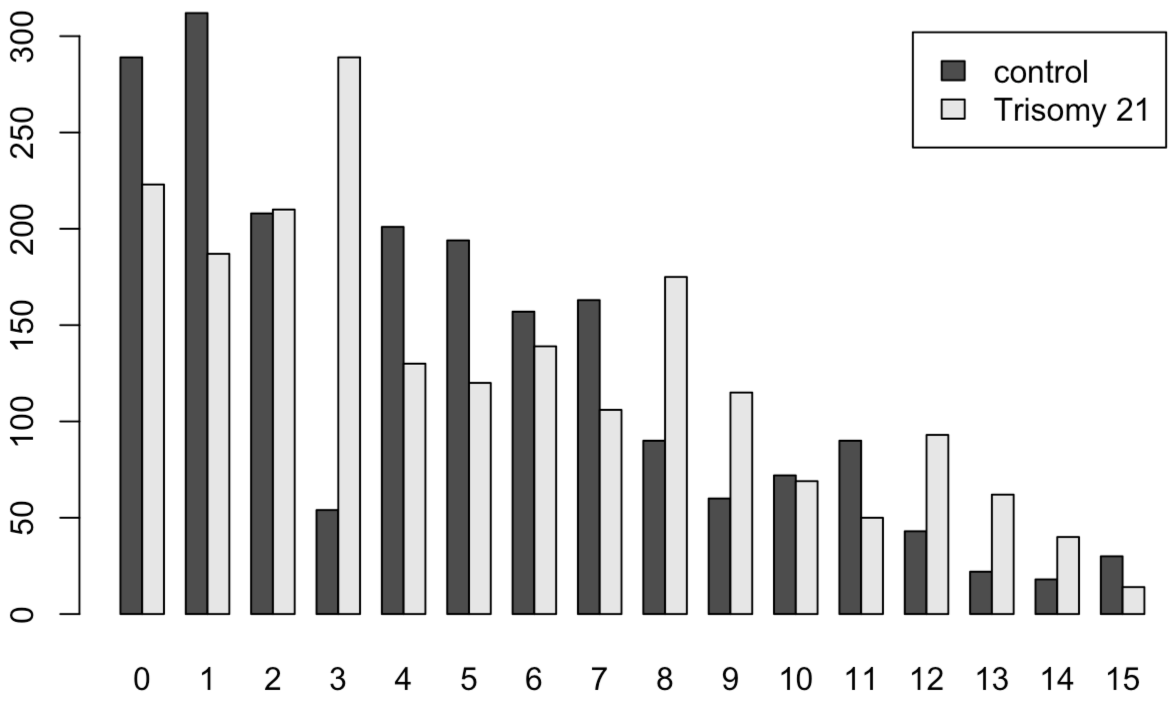


#### 5.2 Fisher's exact test

We use Fisher's exact test to see whether the odds $\frac{\text{trisomy 21 cells in cluster 3}}{\text{trisomy 21 cells in cluster 1}} / \frac{\text{control group cells in cluster 3}}{\text{control group cells in cluster 1}}$ is greater than 1.

```
Testing = matrix(c(312,54,187,289),
                 nrow=2,
                 byrow=TRUE,
                 dimnames = list(Group = c("Control","Trisomy21"),
                                 Cluster = c("Cluster 1","Cluster 3")))
fisher.test(Testing, alternative = "greater")
```

```
##
##  Fisher's Exact Test for Count Data
##
## data:  Testing
## p-value < 2.2e-16
## alternative hypothesis: true odds ratio is greater than 1
## 95 percent confidence interval:
##  6.608068      Inf
## sample estimates:
## odds ratio
##    8.90362
```

Since $p$-value $< 2.2e^{-16} < 0.05$, under $95\%$ confidence level, the odds $\frac{\text{trisomy 21 cells in cluster 3}}{\text{trisomy 21 cells in cluster 1}} / \frac{\text{control group cells in cluster 3}}{\text{control group cells in cluster 1}}$ is significantly greater than 1.

### 6 Detect differentially expressed genes (Positive markers only) for each cluster

#### 6.1 Detect differentially expressed genes

Identify canonical cell type marker genes that are conserved across conditions. We perform differential gene expression testing for each group and combines the p-values using meta-analysis methods from the `MetaDE` R package. We identify differentially expressed genes in each cluster by choosing genes that combined p-values are less than $0.05$.

```
DefaultAssay(combined) <- "RNA"
markers=list()
top5 = c()
for(i in 0:15){
  name <- paste('cluster', i,sep='')
  markers[[name]] <- FindConservedMarkers(combined, ident.1 = i, grouping.var = "grou
p", only.pos = TRUE, verbose = FALSE)
  markers[[name]] <- markers[[name]][(markers[[name]]$max_pval<0.05 &  markers[[nam
e]]$minimump_p_val<0.05),]
  top5=c(top5,rownames(markers[[i+1]])[1:5])
}
```

#### 6.2 Feature plot of PAX6

Finally, we visualize the expression of PAX6 which might be of interest.

```
FeaturePlot(combined, pt.size = 0.3, features = "PAX6", min.cutoff = "q9", split.by =
"group")
```

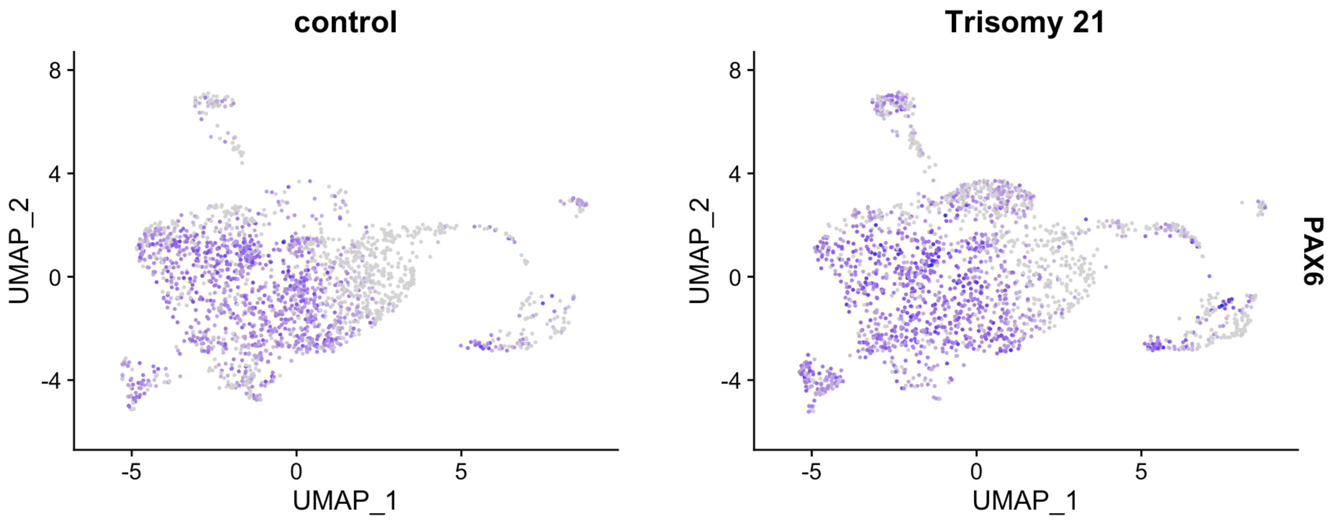


#### 6.3 Additional feature plots

```
FeaturePlot(combined, pt.size = 0.3, features = c("NR2F2","GLI3","NNAT"), min.cutoff
= "q8", split.by = "group")
```

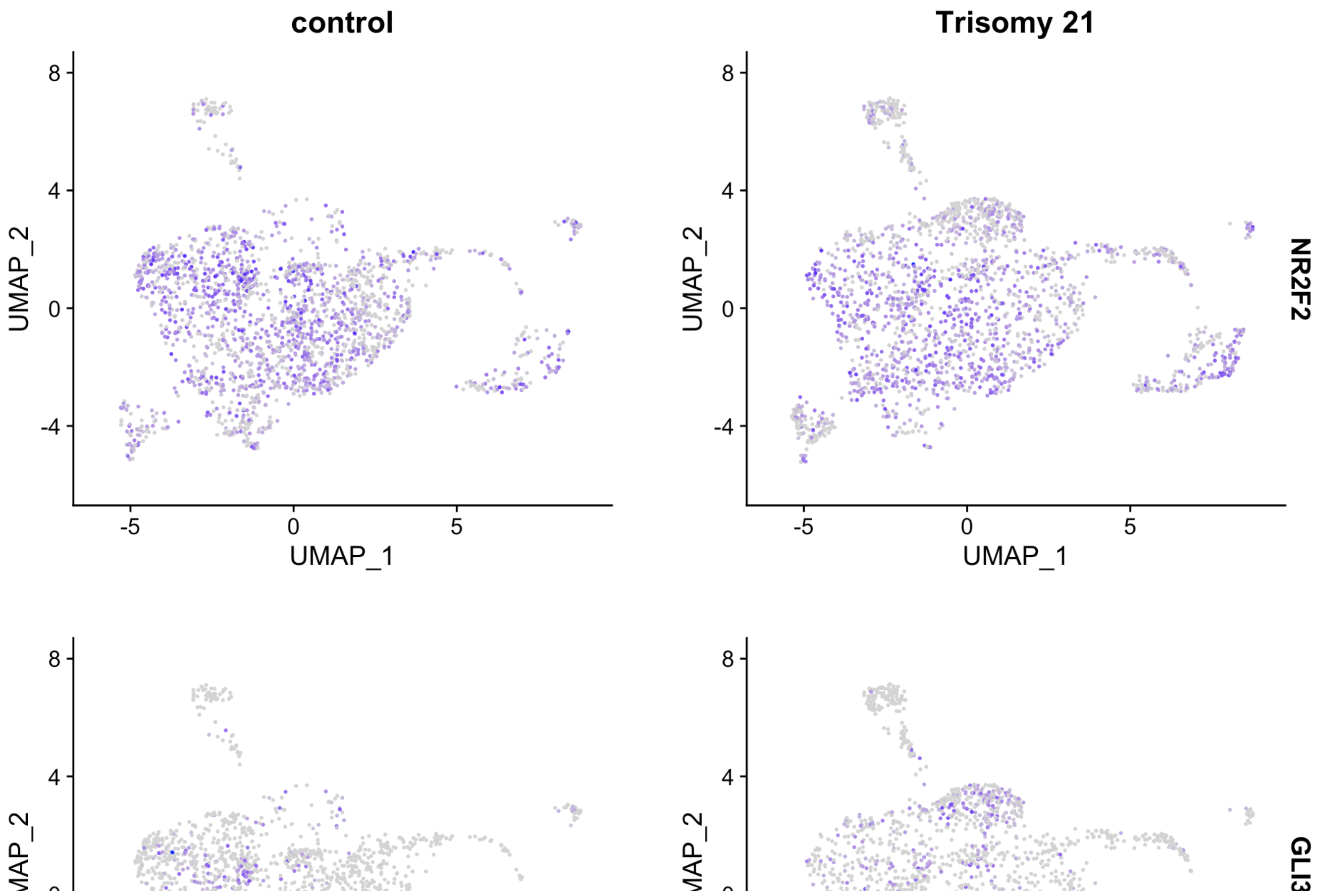

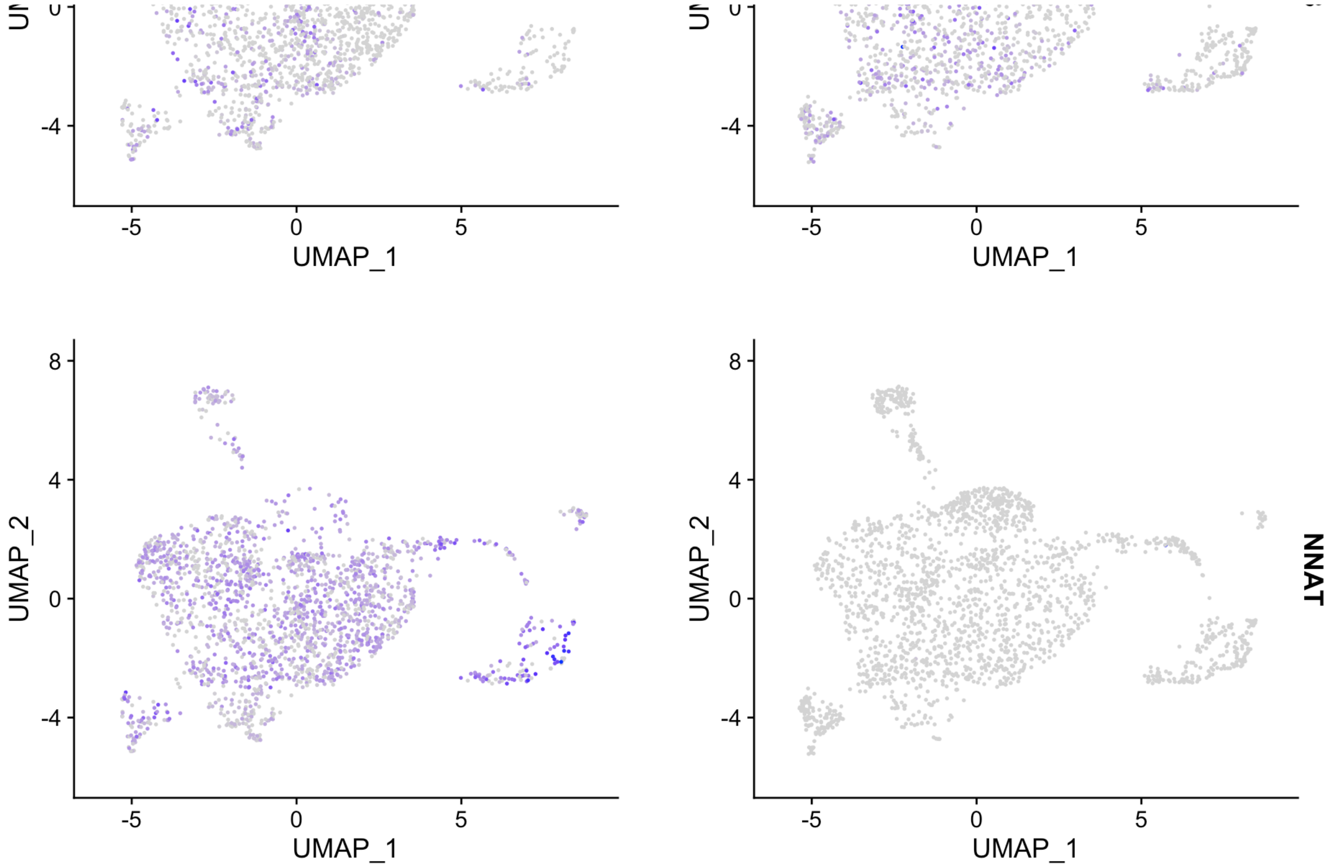


### 7 Detect differentially expressed genes for control vs Trisomy 21

```
dex_genes = FindMarkers(combined,group.by = "group", ident.1 = "control",verbose = FA
LSE)
dex_genes[dex_genes$p_val_adj<0.05,]
```

| | p_val <dbl> | avg_log2FC <dbl> | pct.1 <dbl> | pct.2 <dbl> | p_val_adj <dbl> |
|---|---|---|---|---|---|
| NNAT | 0.000000e+00 | 3.0307762 | 0.799 | 0.001 | 0.000000e+00 |
| METRN | 1.697330e-232 | -0.9467797 | 0.575 | 0.865 | 5.718984e-228 |
| MAGEH1 | 8.006321e-230 | 0.6742327 | 0.455 | 0.018 | 2.697650e-225 |
| ATP5O | 5.622243e-228 | -0.6577688 | 0.966 | 0.996 | 1.894359e-223 |
| POU3F4 | 6.785462e-226 | 0.8315352 | 0.471 | 0.033 | 2.286294e-221 |
| SOD1 | 5.711082e-218 | -0.7263670 | 0.946 | 0.987 | 1.924292e-213 |
| PCSK1N | 2.169855e-188 | -1.0027290 | 0.058 | 0.469 | 7.311109e-184 |
| TTC3 | 3.657587e-150 | -0.6901749 | 0.827 | 0.941 | 1.232387e-145 |

| | p_val | avg_log2FC | pct.1 | pct.2 | p_val_adj |
|---|---|---|---|---|---|
| | <dbl> | <dbl> | <dbl> | <dbl> | <dbl> |
| FGFBP3 | 1.053632e-147 | 1.1088862 | 0.928 | 0.769 | 3.550106e-143 |
| TMSB4X | 5.910802e-137 | 0.6586441 | 1.000 | 1.000 | 1.991586e-132 |

1-10 of 206 rows Previous **1** 2 3 4 5 6 ... 21 Next
